# A consensus variant-to-function score to functionally prioritize variants for disease

**DOI:** 10.1101/2024.11.07.622307

**Authors:** Tabassum Fabiha, Ivy Evergreen, Soumya Kundu, Anusri Pampari, Sergey Abramov, Alexandr Boytsov, Kari Strouse, Katherine Dura, Weixiang Fang, Gaspard Kerner, John Butts, Thahmina Ali, Andreas Gschwind, Kristy Mualim, Jill E. Moore, Zhiping Weng, Jacob C. Ulirsch, Hongkai Ji, Jeff Vierstra, Timothy E. Reddy, Stephen B. Montgomery, Jesse M. Engreitz, Anshul Kundaje, Ryan Tewhey, Alkes L. Price, Kushal K. Dey

**Affiliations:** Computational and Systems Biology, Memorial Sloan Kettering Cancer Center; Department of Genetics, Stanford University, Stanford, CA, USA; Department of Computer Science, Stanford University, Stanford, CA, USA; Altius Institute for Biomedical Sciences, Seattle, WA; Duke Center for Statistical Genetics and Genomics, Duke University, Durham, 27710, NC, USA; Department of Biomedical Engineering, Johns Hopkins University School of Medicine, Baltimore, MD 21205, USA; Department of Biostatistics, Johns Hopkins Bloomberg School of Public Health, Baltimore, MD 21205, USA; Center for Epigenetics, Johns Hopkins University School of Medicine, Baltimore, MD 21205, USA; Department of Epidemiology, Harvard T.H.Chan School of Public Health; The Jackson Laboratory, Bar Harbor, ME, USA; Graduate School of Biomedical Sciences and Engineering, University of Maine, Orono, ME, USA; Basic Sciences and Engineering Initiative, Betty Irene Moore Children’s Heart Center, Lucile Packard Children’s Hospital, Stanford, CA, USA; Department of Plant Biology, Carnegie Institute of Science, Stanford, CA, USA; Program in Bioinformatics and Integrative Biology, University of Massachusetts Chan Medical School, Worcester, MA, USA; Illumina Artificial Intelligence Laboratory, Illumina, San Diego, CA, USA; Division of Integrative Genomics, Department of Biostatistics Bioinformatics, Duke University Medical School, Durham, 27710, NC, USA; Program in Computational Biology Bioinformatics, Duke University, Durham, 27710, NC, USA; Department of Pathology, Stanford University, Stanford, CA, USA; Novo Nordisk Foundation Center for Genomic Mechanisms of Disease, Broad Institute, Cambridge, MA, USA; Stanford Cardiovascular Institute, Stanford University, Stanford, CA, USA; Broad Institute of MIT and Harvard, Cambridge, MA, USA; Department of Biostatistics, Harvard T. H. Chan School of Public Health; Physiology, Biophysics and Systems Biology, Weill Cornell Medicine; Gerstner Sloan Kettering Graduate School of Biomedical Sciences

## Abstract

Identifying and functionally characterizing causal disease variants in genome-wide association studies remains a pressing challenge. Here, we construct a consensus variant-to-function (cV2F) score that assigns a single value to each common single-nucleotide variant in the genome, and helps to predict and characterize causal disease variants. The cV2F score leverages features reflecting variant-level experimentally and computationally predicted function (e.g. allelic imbalance and sequence-based deep learning models) and element-level function (e.g. predicted enhancers), and learns optimal combinations of features by training a gradient boosting model on GWAS fine-mapping results. The cV2F-annotated variants attained an AUPRC of 0.822 at identifying held-out fine-mapped variants. Variants with high cV2F scores are highly enriched for heritability (14.2x, s.e. 0.5) across 66 diseases/traits, are uniquely informative for disease heritability, and are highly predictive of variants implicated by reporter assays; cV2F substantially outperforms previous variant-to-function scores using all of these metrics. GWAS fine-mapping of 110 diseases/traits informed by cV2F identified 14.3% more confidently fine-mapped (PIP > 0.95) variants than non-functionally informed fine-mapping. We further constructed tissue/cell line-specific cV2F scores that prioritize variants based on regulatory potential in specific tissues/cell lines, attaining high heritability enrichment for tissue-related diseases/traits (15.6x, s.e. 2.3) while providing independent information (average correlation of 0.27 with the primary cV2F score). We highlight examples of GWAS loci for which cV2F pinpoints causal variants with high confidence and elucidates their functional role.

## Introduction

Genome-wide association studies (GWAS) have significantly advanced the identification of disease-associated variants^1–3^, but fine-mapping these associations to pinpoint causal variants remains challenging due to extensive linkage disequilibrium between variants^4–8^. Functional genomics data encompassing various biochemical assays^9–12^, functional characterization experiments^13–16^, and sequence-based computational models^17–22^ provide a wealth of information complementary to GWAS regarding variant function. Developing strategies to combine this functional information with GWAS to enhance the identification of disease-causal variants is of utmost importance, mirroring integrative approaches that prioritize disease genes^23^ and variants for shared common and Mendelian disease risk^24^. Current integrative pathogenicity scores are limited to the diversity of assays and resolution of functional information used, are not informed by GWAS data, and/or are aimed at rare Mendelian variants^25,26^.

Here, we propose a new consensus variant-to-function (cV2F) score that aggregates evidence of biological function across a broad set of functional features reflecting variant-level and element-level function, training a gradient boosting model on GWAS fine-mapping results to learn optimal combinations of features. We show that cV2F outperforms previously proposed scores across a range of metrics spanning held-out fine-mapping data, functional components of disease heritability, and experimentally validated variants. We show that incorporating the cV2F score in GWAS fine-mapping increases the number of confidently fine-mapped variants. We further show that tissue/cell line-specific cV2F scores restricted to features from focal tissues/cell lines are highly disease informative and provide independent information.

## Results

### Overview of methods

The consensus Variant-to-Function (cV2F) score assigns a binary 0/1 value to each variant based on evidence of variant-level and element-level regulatory function. The cV2F score is generated using a non-linear combination of variant-level and element-level regulatory features from the ENCODE consortium (Phase 4), baseline-LD model (v 2.2) features commonly used in disease heritability analyses^27,28^ and additional conservation features^29,30^ (**Figure 1A, Supplementary Table S1**). The optimal non-linear combination is determined by a gradient-boosting classification model^31,32^ trained on GWAS fine-mapped variants (Methods, Figure 1B). This model generates a probabilistic grade (between 0 and 1) to each variant, which we term as “probabilistic cV2F score”; this probabilistic score is binarized (score > 0.75) to generate a binarized score, which we call the “cV2F score”. The cV2F score is computed for 10 million SNPs with minor allele count >=5 in 1000 Genomes Project Europeans^33^, analogous to previous work^28^.

**Figure 1:**
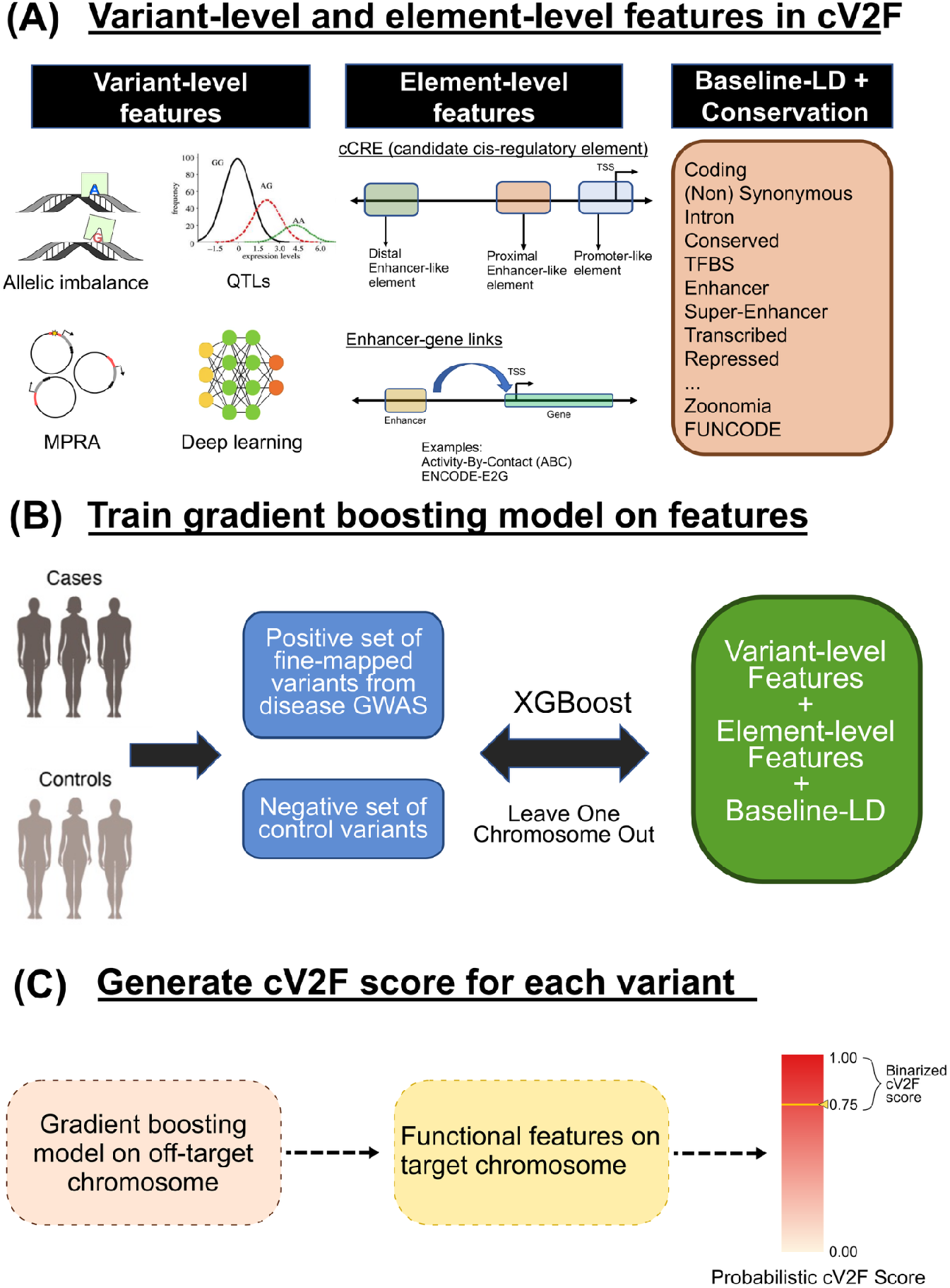
Consensus Variant-to-Function (cV2F) Score. (A) An illustration of variant-level regulatory function, element-level regulatory function and baseline-LD annotations that are used as features in the cV2F gradient boosting model. (B) The cV2F XGBoost gradient boosting^32^ based classification model using GWAS fine-mapped informed variants as positive and negative labeled sets. (C) A schematic of how for a specific variant, the functional features at the variant are optimally aggregated using the gradient boosting model from (B) fitted on all other chromosomes, to generate the cV2F probabilistic grade for the variant.

The cV2F probability model is of the form

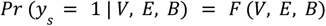

where *y*_*s*_ is a binary indicator (1 or 0) indicating whether the variant *s* is a disease-causal variant based on standard statistical fine-mapping, *V, E* and *B* denote a total of 339 variant-level, element-level and baseline-LD features, and *F* is a non-linear function of the features that is most predictive of *y*_*s*_; *F* is optimized by training an XGBoost gradient boosting model^31,32^ on a set of positive and negative control variants (*y*) derived from GWAS fine-mapping using the 339 features (**Figure 1B**). For positive control variants, we used variants that were confidently fine-mapped^5^ (PIP > 0.90) for one or more of 94 UK Biobank GWAS diseases and traits^34^; for each positive control variant, we selected 5 negative control variants in the same LD block with the most similar MAF that had low PIP (< 0.01) across all 94 UK Biobank traits (**Methods**). We considered other positive and negative control sets informed by allelic effect results from a large-scale MPRA experiment^13^ and WG-STARR-seq assay (T. Reddy, unpublished data, **Data Availability**), variants with significant regulatory impact from 3 aggregated CRISPRi experiments^35–37^, and fine-mapped eQTLs from GTEx^38^. To avoid overfitting, cV2F scores are trained using a leave-one-chromosome-out framework (**Figure 1C**). Tissue/cell line-specific cV2F scores are generated by restricting features to variant-level and element-level features from a selected set of tissues/cell lines (see **Supplementary Table S1**). We have publicly released open-source software implementing cV2F (**Code Availability**).

The 339 features include 186 variant-level features reflecting different modalities such as allelic imbalance calls in genomic DNase I footprinting^12^ (J. Vierstra, unpublished data, **Data Availability**), TF ChIP-seq data^39^, allele-specific effects in reporter assays^13^, predicted allele-specific effects from 3 deep learning models (Enformer^21^, ChromBPNet^40,41^ (A. Kundaje, unpublished data, **Data Availability**), an MPRA data-driven deep learning model (R. Tewhey, unpublished data, **Data Availability**), and fine-mapped eQTLs^42^; 52 element-level features derived from enhancer and promoter signatures in the updated candidate cis-regulatory elements (cCRE)^10^ (version 4) (J. Moore, Z. Weng unpublished data, **Data Availability**) and enhancer predictions from the Activity-By-Contact (ABC) model^11,34,35^; 84 features based on variant-level and element-level coding, epigenomic, conservation annotations from the baseline-LD (v2.2)^27,43^ model (MAF bin and LD annotations were omitted as the training adjusts for MAF and LD differences between positive and negative control variants); and 17 features based on additional conservation annotations from the FUNCODE^44^ (**Data Availability**) and Zoonomia projects^29,30^ (**Supplementary Table S1**). For each feature, we annotated 9,991,229 variants (hg38) with minor allele count >=5 in 1000G Europeans^33^. For features with missing data for some variants, we set values of those variants to the mean value; we show below that this choice did not impact our predictive accuracy. We restricted our focus to features aggregated across 9 different tissues (blood, brain, kidney, liver, heart, lung, intestine, fat, skin) inferred to be critical for some complex diseases and traits^45^, and 5 ENCODE cell-lines (K562, GM12878, HepG2, SKNSH and A549).

We evaluated cV2F scores in three ways. First, we computed the area under the precision and recall curve (AUPRC) on leave-one-chromosome-out sets of positive control and negative control variants. Second, we computed excess overlap and recall (**Methods**) with respect to (i) variants with significant allelic effects in an MPRA dataset targeting GWAS-implicated variants^13^ and (ii) a WG-STARR-seq dataset targeting African genetic variation in K562 (T. Reddy, unpublished data, **Data Availability**); we conservatively remove all reporter assay features from cV2F when computing this metric to avoid any circularity. Third, we computed two metrics, heritability enrichment and standardized effect size (***τ***\*), to score annotations in the S-LDSC disease heritability framework^28^, applied also in a leave-one-chromosome-out fashion on the cV2F score, similar to ref^46^. Heritability enrichment is defined as the proportion of heritability explained by SNPs in an annotation divided by the proportion of SNPs in the annotation. Standardized effect size (***τ***\*) is defined as the proportionate change in per-SNP heritability associated with a 1 standard deviation increase in the value of the annotation, conditional on other annotations included in the model^47^. Unlike heritability enrichment, ***τ***\* quantifies effects that are unique to the focal annotation, by conditioning on other annotations; thus, we use ***τ***\* as our primary metric.

We used the primary and tissue-specific cV2F scores to perform functionally informed fine-mapping by reweighting the posterior probability for each credible set obtained from non-functionally informed fine-mapping^5^, analogous to ref.^38^ (**Methods**). We have publicly released primary and tissue-specific cV2F scores computed in this study, as well as functionally informed fine-mapping results for 110 UK Biobank diseases and traits (**Data Availability)**.

### Constructing a consensus Variant-to-Function score

We fit the cV2F gradient boosting model for the full set of 339 features, as well as different subsets of features. The highest predictive accuracy of the classification task was attained using all variant-level, element-level, baseline-LD and other conservation features (AUPRC=0.822) (**Figure 2A, Supplementary Table S2**). The addition of variant-level, element-level and other conservation features on top of the baseline-LD features resulted in a significantly higher AUPRC (p-value = 0.006, 2-sample t-test; 13.5% improvement in 1−AUPRC). The variant-level features attained a significantly higher AUPRC vs. element-level features (0.777 vs. 0.669; p-value=2e-06), highlighting the importance of assessing function at base pair resolution. The element-level features did not attain a significant improvement to AUPRC when added to variant-level features (0.782 vs. 0.777; p-value = 0.54). We note that element annotations form the backbone for most sequence models of variant function and can be easily deployed more broadly across cell types and tissues compared to functional experiments like MPRA or CRISPR. Among the variant-level features, sequence-based deep learning model features attained the highest AUPRC (0.725), although the addition of other experimentally and genetically informed variant-level features led to a significant increase in predictive accuracy (23.3% improvement in 1-AUPRC, p=3e-06) (**Figure 2A)**. We note that the model performance is dependent on the number of biosamples, epigenomic features, and number of variants tested by each model or experimental assay. For example, experimental features have lower predictive accuracy compared to deep learning models likely owing to the limited number of variants tested (1.8% for MPRA and 12% for DNase allelic imbalance). Additionally, restricting Enformer features to DNase (and excluding TF ChIP-seq, Histone modification ChIP-seq and CAGE feature tracks) produced more comparable predictive performance as ChromBPNet model trained on DNase data (0.683 versus 0.670; p-value = 2e-02) (**Supplementary Figure S1A**). Baseline-LD features attained higher AUPRC (and produced a larger drop in predictive performance upon ablation) compared to ENCODE variant-level and element-level features; this is expected as baseline-LD features span broader functional categories involving coding, conserved, and LD-related annotations while ENCODE annotations are regulatory in nature (**Figure 2A, Supplementary Figure S1B**). We performed an additional analysis by training the model using single features from the cV2F training model and reporting the AUPRC for each model; this provides an assessment of the relative importance of each feature marginally in predicting GWAS fine-mapped variants. Enformer and conservation-related annotations like PhastCons^48^ and GERP-NS^49^ showed the highest predictive accuracy (**Supplementary Figure S1C**). We reached similar conclusions about feature importance based on the Shapley values^50^ of these functional annotations (**Supplementary Figure S1D**).

**Figure 2:**
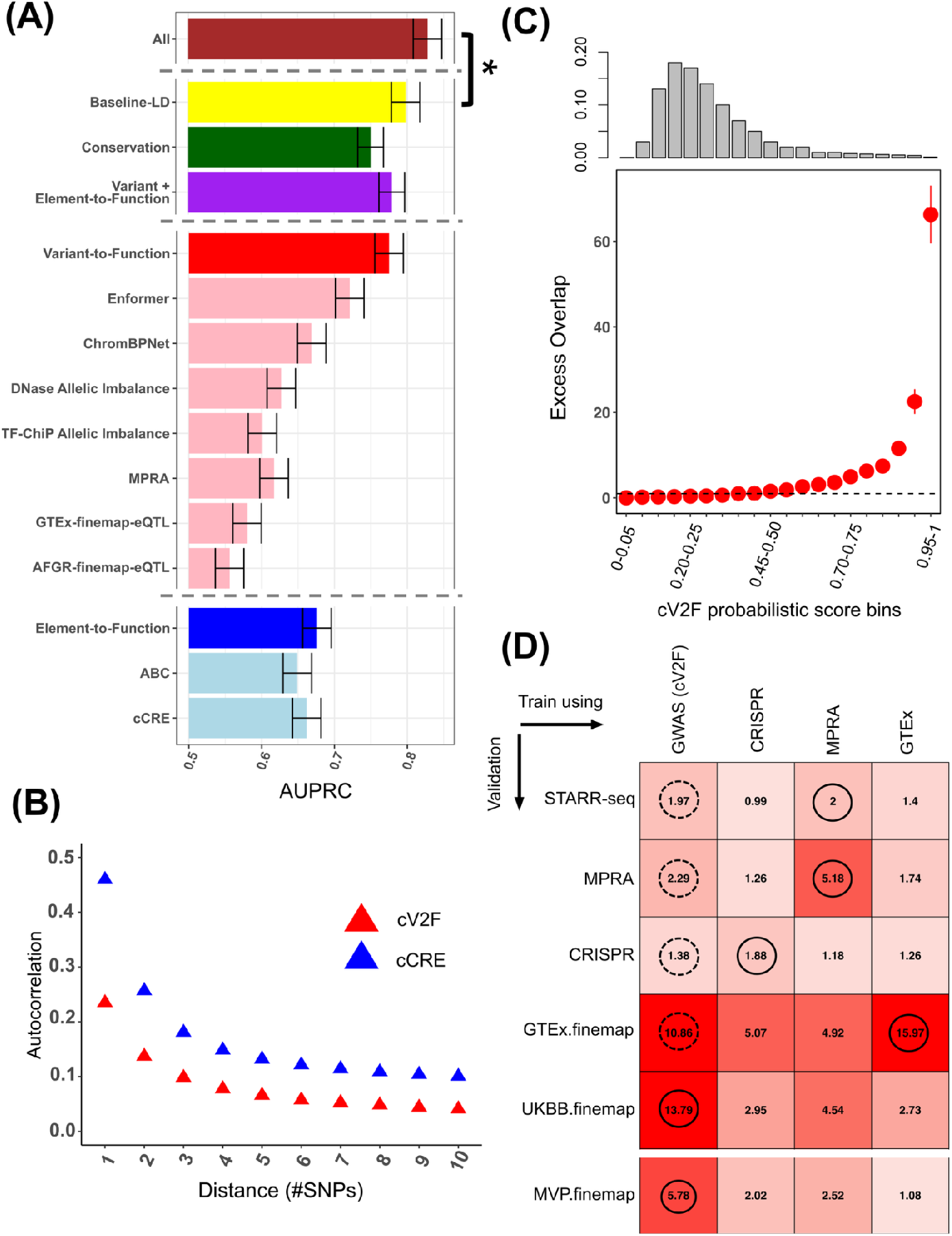
Properties of the consensus Variant-to-Function (cV2F) Score. (A) Area under the Precision Recall curve (AUPRC) of the cV2F gradient boosting model on held-out sets of GWAS positive (PIP> 0.90 variants in at least one out of 94 UKBB diseases and traits) and LD and MAF matched negative variants (PIP < 0.01 across all traits), using different sets of functional features. These functional features can be broadly grouped into three categories - baseline-LD (yellow), conservation scores from Zoonomia and FUNCODE efforts (green) and ENCODE variant and element-level functional annotations (purple). We further report the AUPRCs for categories of features (pink) that make up the ENCODE variant-function annotations (red) and categories of features (lightblue) that make up the ENCODE element-function annotations (blue). Error bars denote 95% confidence intervals. (B) The autocorrelation in the ENCODE cCRE and primary binarized cV2F score for variants that are between 1st and 10th nearest in physical distance to each variant in the genome. (C) Excess overlap of variants in different bins of cV2F scores with respect to variants confidently fine-mapped with PIP > 0.75 for one or more of 94 UKBiobank GWAS traits. The histogram of the number of variants in each bin of cV2F scores is shown on the top. (D) Enrichment of the top 2.4% variants (matched size with cV2F > 0.75) identified by training the same set of features on 4 different training datasets derived from GWAS fine-mapping data from 94 UKBB traits for cV2F, CRISPR screens in K562, MPRA data in 5 cell lines and fine-mapped eQTL in 49 GTEx tissues. The bold circle denotes the top-performing method (in column) for each row, while the dotted circle denotes the second-best method. Numerical results are reported in **Supplementary Table S2**.

We computed the genomic autocorrelation of the cV2F probabilistic score (i.e. the correlation between consecutive variants) to assess its effectiveness in precise genomic localization of variants with predicted function. The cV2F probabilistic score exhibited a 2.3x lower autocorrelation compared to the ENCODE element-level cCRE annotations (**Figure 2B, Supplementary Table S2**). We compared the autocorrelation of the probabilistic cV2F score against probabilistic scores trained on subset of element and variant level features from **Figure 1A**; as expected, the proposed probabilistic cV2F score exhibited 2.7x lower autocorrelation than the analogous probabilistic score trained only on element-level features, highlighting the importance of variant-level features (**Supplementary Figure S2A**).

We assessed the enrichment of different bins of cV2F probability grades in held-out sets of fine-mapped SNPs. We observed a non-linear distribution in this enrichment across cV2F bins (**Figure 2C)**; this may be attributed to the fact that the cV2F training model uses positive and negative sets of variants that are putatively more and less functionally important than an average SNP in the genome respectively, which leads to an average SNP in the genome having a moderate value cV2F score (mean cV2F score = 0.29 across the genome). We assessed the enrichment and recall of variants implicated by binarized cV2F at different binarization thresholds with respect to (i) weakly fine-mapped SNPs (PIP > 0.10) recommended for benchmarking in ref.^34^, and (ii) MPRA and STARR-seq positives versus tested variants based on allelic effects; for both these comparisons, cV2F scores thresholded at 0.75 performed the best in terms of both precision and recall (**Supplementary Figure S2B-D**). The binarized cV2F score thresholded at 0.75 identified 237,600 variants (2.4% of all variants) as important for disease-associated function (**Supplementary Table S3**, link in **Data Availability**). We observed a high 4.9x (s.e. 0.04x) excess overlap of binarized cV2F variants with variants in enhancer-annotated functional regions^51^, as well as high excess overlap in other functional regions such as promoter, coding, TSS and UTRs (excess overlap between 4.3x-14.1x) (**Supplementary Figure S2E**); thus cV2F is effective in capturing enhancer-driven disease-critical regulatory architecture besides that of other functional regions. Variants annotated by binarized cV2F have only slightly lower minor allele frequency (MAF) compared to an average variant in the genome; suggesting that MAF differences are not a driver of the cV2F score (**Supplementary Table S2)**.

To assess the effectiveness of using GWAS fine-mapped variants as the training data for cV2F, we compared the cV2F score to scores generated using the same set of features but 3 other training data sets of positive control (resp. negative control) sets of variants. These included variants with significant allelic effects (resp. no allelic effect changes) in large-scale MPRA^13^ and multiple CRISPRi screening assays^35–37^, and variants that are fine-mapped eQTLs in any GTEx tissue^38^ (resp. not fine-mapped eQTLs in any GTEx tissue). When using GTEx eQTL for training, both AFGR QTL, GTEx eQTL and MPRA results on GTEX-selected variants were dropped from the feature set to avoid circularity. When using MPRA data for training, all reporter assay features were dropped from the feature set to avoid circularity. We note that the CRISPR data were not included as features due to the small number of features. We assessed the enrichment of the top 2.4% of scored variants (same size as binarized cV2F) in 6 different validation sets - MPRA and STARR-seq (also used for cV2F validation below), CRISPR positives, confidently fine-mapped (PIP > 0.90) GWAS variants from UKBiobank^52^ and Million Veteran Program (MVP)^53^, and confidently fine-mapped (PIP > 0.50) GTEx eQTLs. All scores were trained and validated in a leave-one-chromosome-out framework. Results are reported in **Figure 2D** and **Supplementary Table S2**. For a given validation set we expectedly observed that the training data set most similar to that validation set performed best, but the cV2F score trained on GWAS data consistently performed among the top two methods across different validation sets. The cV2F score also showed strongest enrichment (compared to other methods) in the held-out cohort of MVP multi-ancestry fine-mapped variants across 931 GWAS traits, implying that cV2F-implicated variants can be used to identify functional variants for traits spanning multiple ancestries. The alternative training data sets had lower predictive accuracy (AUPRC) vs. the GWAS fine-mapping training data, however, the full combination of variant-level, element-level, baseline-LD and other conservation features continued to perform best (**Supplementary Figure S2F**).

We performed 5 secondary analyses. First, we trained models with the same feature sets using positive and negative sets of fine-mapped variants from the Million Veteran Program^53^(MVP; European-American or African-American GWAS data for 931 traits) instead of the UK Biobank; we continued to use positive and negative sets of fine-mapped variants from the UK Biobank for validation. Results for both European-American and African-American data were similar to **Figure 2A** in both absolute and relative accuracies for different feature sets (**Supplementary Figure S3A,B**); accuracies were similar when using MVP data for validation (**Supplementary Figure S3C**). These results confirm that functional architectures of disease are consistent across ancestries^54–56^. Second, we checked that setting feature values of variants with missing data in the experimental assays to the mean value did not impact our predictive accuracy. Specifically, we verified that restricting training and validation data to variants with non-missing values for experimental features testing a subset of variants did not impact prediction accuracy (**Supplementary Table S2**). Third, we checked that restricting the deep learning annotations from Enformer and ChromBPNet to the set of tested variants from DNase allelic imbalance experimental assay, and mean-imputing the remaining variants results in considerable drop in predictive accuracy; this underscores that the number of variants tested can impact the overall predictive accuracy of features (**Supplementary Table S2**). Fourth, we evaluated the prediction accuracy of cV2F when trained on lower PIP thresholds of fine-mapping in order to accommodate more variants in its positive training set; we observed significantly reduced performance when using low-confidence fine-mapped variants for training (**Supplementary Table S2**).Fifth, in order to assess the impact of conservation on the overall predictive accuracy of element-level and variant-level functional features, we split the training data into high and low conservation bins using GERP and Zoonomia (PhastCons) scores. Across all conservation bins, variant-level features consistently outperformed element-level features; however the overall predictive accuracy was notably lower in bins with lower conservation scores **(Supplementary Figure S3D,E)**.

We conclude that the cV2F score provides an optimal combination of functional features across different assays and predictive models and attains high enrichment across functional and genetic characterizations.

### The consensus Variant-to-Function score is informative for disease

We assessed the (binarized) cV2F score using the S-LDSC disease heritability framework^28,57^. We compared the heritability enrichment and standardized effect size (***τ***\*) conditional on the baseline-LD model (v2.2)^43^ of the cV2F score to 5 previously proposed pathogenicity scores: Regulomedb^58,59^, MACIE^60^, CADD^25,26^, EigenPC^61^ and ReMM^62,63^ (**Methods**). We meta-analyzed results across 10 brain-related traits (brain), 15 blood-related traits (blood), or a broad set of 66 diseases/traits (all) analyzed in ref^45^ (**Supplementary Table S1**). Results are reported in **Figure 3A** and **Supplementary Table S4**. In both the all and blood meta-analyses, cV2F attained comparable or higher enrichments (average 14.2x, se 0.53) than all other scores (including those with much smaller annotation sizes: CADD and ReMM), and much higher ***τ***\* values (≈2) than all other scores; annotations with ***τ***\* value >0.5 are generally considered to be highly important for disease^64^. In the brain meta-analysis, ***τ***\* values were non-significant after Bonferroni correction for all scores, consistent with the known phenomenon of weaker functional enrichment for brain-related traits^4,39,40^. The binarized cV2F score also consistently outperformed the probabilistic cV2F score and other versions of Regulomedb and MACIE scores (**Supplementary Figure S4, Supplementary Table S4**). cV2F also outperformed a broader set of 10 other Mendelian disease-derived pathogenicity scores^65–77^ (**Supplementary Figure S5A-B, Supplementary Table S5**); boosted versions of all other pathogenicity scores, boosted using AnnotBoost^24^ (**Supplementary Figure S5C-D, Supplementary Table S5**); and all other pathogenicity scores when analyzed jointly using stepwise elimination to iteratively remove conditionally non-significant annotations (**Supplementary Figure S6A, Supplementary Table S5**).

**Figure 3:**
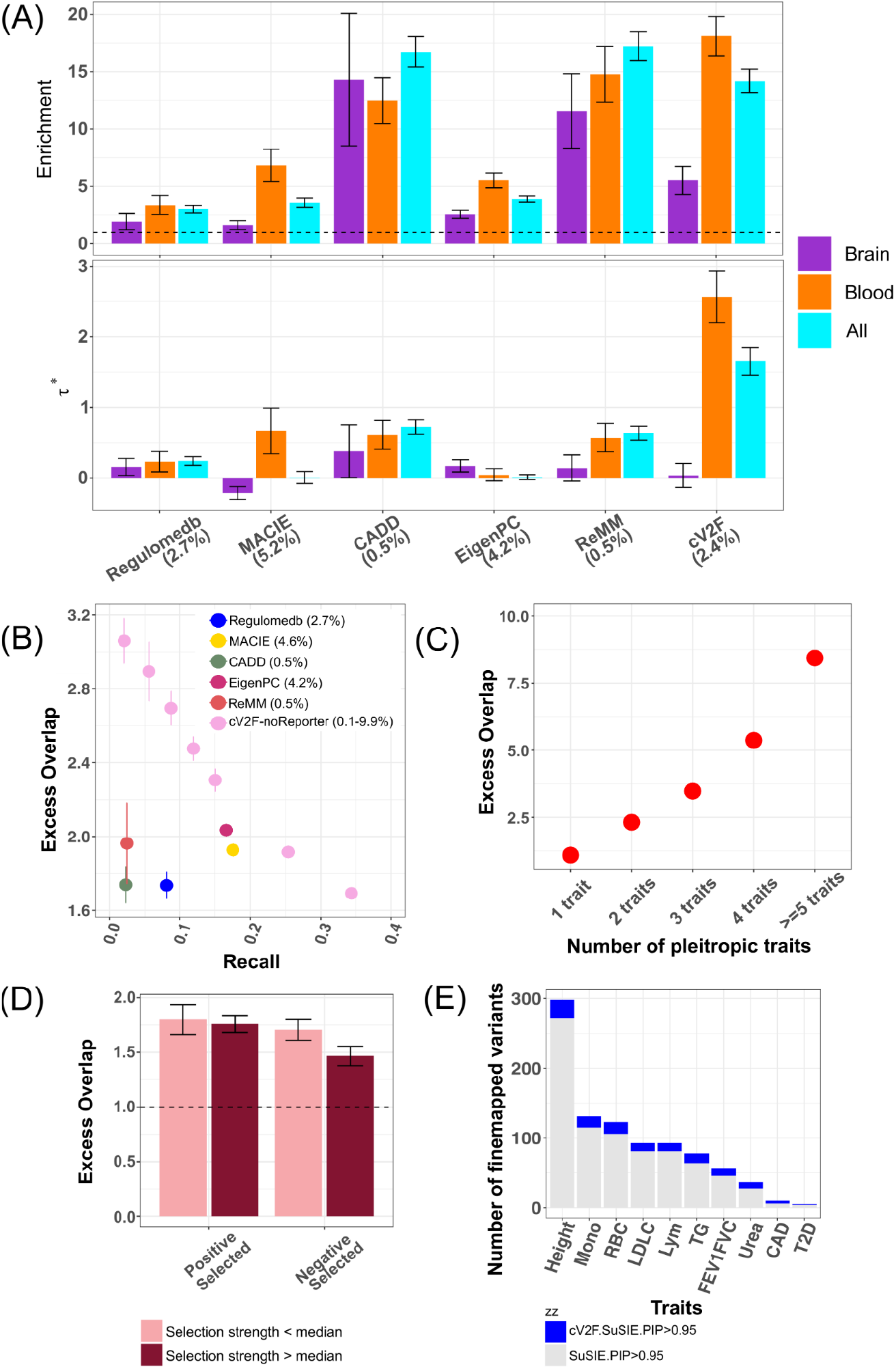
Benchmarking the cV2F score against other approaches. (A) S-LDSC Heritability enrichment (top panel) and standardized effect sizes (τ*) (bottom panel) of optimally binarized cV2F score at threshold of 0.75, Regulomedb annotated variants (binarized at probability > 0.90), MACIE-anyclass annotated variants (binarized at threshold of 0.99), and 3 primary genome-wide and near genome-wide Mendelian disease-derived pathogenicity scores (CADD^25,26^, Eigen-PC^65^ and ReMM^62,63^) optimally binarized based on the recommendation from ref^24^. See **Supplementary Figures S4, S5** for similar heritability based benchmarking against other approaches. Dashed horizontal line denotes no enrichment. Results are meta-analyzed across all 66 relatively independent diseases and traits, as well as 15 relatively independent blood-related traits and 10 relatively independent brain-related traits. (B) Excess overlap and recall of variants annotated by different variant-function and pathogenicity scores with respect to variants that are MPRA positives among the ones tested for functional characterization, in comparison to the cV2F-noReporter score (an analogous score to cV2F but not including any MPRA related features), binarized at different thresholds. (C) Excess overlap of variants annotated by binarized cV2F in variants that are pleiotropically fine-mapped in 1 to 5 relatively genetically uncorrelated (rg < 0.8) traits. (D) Excess overlap of variants annotated by binarized cV2F in variants that are candidate positively selected variants with selection strength less than and more than the median, and candidate negatively selected variants with selection strength less than and more than the median, where the positive and negatively selected variants are determined based on adaptation to pathogens as in ref^78^. (E) Number of variants confidently fine-mapped (posterior probability of causality > 0.95) from standard fine-mapping and cV2F-informed functional fine-mapping of 10 selected diseases and traits covering the overall spectrum of relative improvement in fine-mapping across all 94 UKBB traits. See **Supplementary Figure 5D** for the full set of traits. Error bars denote 95% confidence intervals. Numerical results are reported in **Supplementary Table S4**.

We assessed the performance of cV2F with respect to variants with significant allelic effects in an MPRA dataset targeting GWAS-implicated variants^13^ and a WG-STARR-seq dataset targeting African genetic variation in K562 (T. Reddy, unpublished data, **Data Availability**); to avoid circularity, we conservatively removed reporter assay features from cV2F in this analysis (cV2F-noReporter). We assessed the excess overlap and recall of the cV2F-noReporter score binarized at different thresholds with respect to the union of all positive variants (versus all tested variants) in MPRA experiments (targeting 136,173 variants at GWAS-implicated loci from UKBiobank^52^ and Biobank Japan^56^, in 5 cell lines) or WG-STARR-seq experiments (targeting 204,802 variants in K562). Analogous to above, we compared cV2F-noReporter to the 5 previously proposed variant-function scores from **Figure 3A**. We determined that cV2F-noReporter binarized at different thresholds attained higher excess overlap and/or recall than each of the other methods (**Figure 3B, Supplementary Figure S6B**, and **Supplementary Table S4**).

Next, we evaluated whether the cV2F score more strongly prioritizes variants that impact multiple diseases and traits. To this end, we assessed the excess overlap of the (binarized) cV2F variants with variants that were suggestively fine-mapped (PIP > 0.10; recommended for benchmarking in ref^34^) for either one trait or multiple traits (restricting to 79 relatively genetically uncorrelated traits (*r*_*g*_ < 0.8) (**Figure 3C, Supplementary Table S4)**. Variants that were pleiotropically fine-mapped for multiple traits exhibited a clear pattern of higher excess overlap with cV2F variants, e.g. excess overlap of 8.4 (s.e. 0.12) for variants fine-mapped for 5 or more traits. We also assessed the excess overlap of the cV2F variants with a set of 21,129 candidate positively selected and 24,152 candidate negatively selected variants based on genetic adaptation to selective pressure exerted by pathogens^78^ (**Figure 3D, Supplementary Figure S6C, Supplementary Table S4**). cV2F variants exhibited highly significant excess overlap for both positively selected and negatively selected variants; the excess overlap did not increase with the strength of selection.

We used the probabilistic cV2F score to perform functionally informed fine-mapping of variants associated with 110 UK Biobank diseases/traits (**Methods**) by re-weighting posterior probabilities of non-functionally informed SuSIE^5^ fine-mapping credible sets, analogous to ref^16^ (**Data Availability**). We observed a 14.3% increase in the number of confidently fine-mapped (PIP > 0.95) variants when incorporating the cV2F score (3,551 vs. 3,107 PIP > 0.95 variants), with a median improvement of 15.3% across traits; the relative improvement was larger for less well-powered traits (**Figure 3E, Supplementary Figure S6D** and **Supplementary Table S6**). We also observed a 20.6% reduction in the aggregate sizes of 95% credible sets across all traits (**Supplementary Table S6**). cV2F-informed fine-mapping detected a larger number (3551 versus 3,438) of confidently fine-mapped (PIP>0.95) variants compared to fine-mapping using functionally informed prior causal probabilities from the baseline-LD model^27^ (with meta-analyzed enrichments from real data analyses), as implemented in PolyFun+SuSIE^6^ (14.3% improvement vs. 10.7% improvement compared to non-functionally informed fine-mapping; **Supplementary Table S6)** (see **Discussion**). We note that the cV2F score prioritizes variants based on pleiotropic function across multiple diseases; as a consequence, fine-mapping that relies on cV2F may overlook variants that have highly specific regulatory functions pertinent to particular diseases.

Linking prioritized variants to the genes that they regulate is a fundamental goal^11,34^. We sought to assess if genes linked to fine-mapped variants prioritized by the cV2F score are more relevant to disease than genes linked to all fine-mapped variants. We analyzed 35,237 suggestively fine-mapped variants (PIP > 0.10) for 94 UKBiobank traits^34^, of which 4,867 (13.8%) were prioritized by the cV2F score. We linked these variants to genes using the cS2G strategy^79^, implicating 1,148 genes using fine-mapped variants prioritized by the cV2F score vs. 2,155 genes using all fine-mapped variants. The 1,148 genes had a 1.18x (p=1.7e-36) higher PoPS prioritization score^23^ for the corresponding traits than the full set of 2,155 genes (**Supplementary Figure S7A**), implying that genes implicated using cV2F are more relevant to disease. We also observed significant excess overlap of these genes in approved drug targets^80^, genes linked to Mendelian disorders^81^ and Enhancer Domain Score (EDS)^82^ implicated genes (**Supplementary Figure S7B**).

We conclude that the cV2F score contains unique information about disease, and can be used to improve fine-mapping of causal disease variants.

### Tissue-specific cV2F provides unique information about disease

We generated tissue/cell line-specific cV2F scores for 3 tissues (blood, brain, liver) and 3 ENCODE cell lines (K562, GM12878, HepG2) for which there are >5 well-matched relatively independent heritable diseases and traits. We restricted the cV2F model to functional features corresponding to ENCODE variant-level and element-level features assayed in the focal tissue/cell line; we used the full set of 94 UK Biobank traits for training. We observed moderately high classification accuracy for all tissues/cell lines, with AUPRC ranging from 0.68-0.72 (**Figure 4A top panel, Supplementary Table S2**); this is lower than the AUPRC of 0.822 for the primary cV2F model (because non-tissue-specific features were not included). Results of specific subsets of tissue-specific features for one tissue (liver) are reported in **Supplementary Table 2**. We binarized each tissue/cell line-specific cV2F score using the same threshold of 0.75 that we used for the primary cV2F score, producing binarized scores of similar size (1.3-2.4% of variants, vs. 2.4% of variants for primary cV2F score) but with low correlation to the primary cV2F score (average 0.27; **Supplementary Figure S8A**).

**Figure 4:**
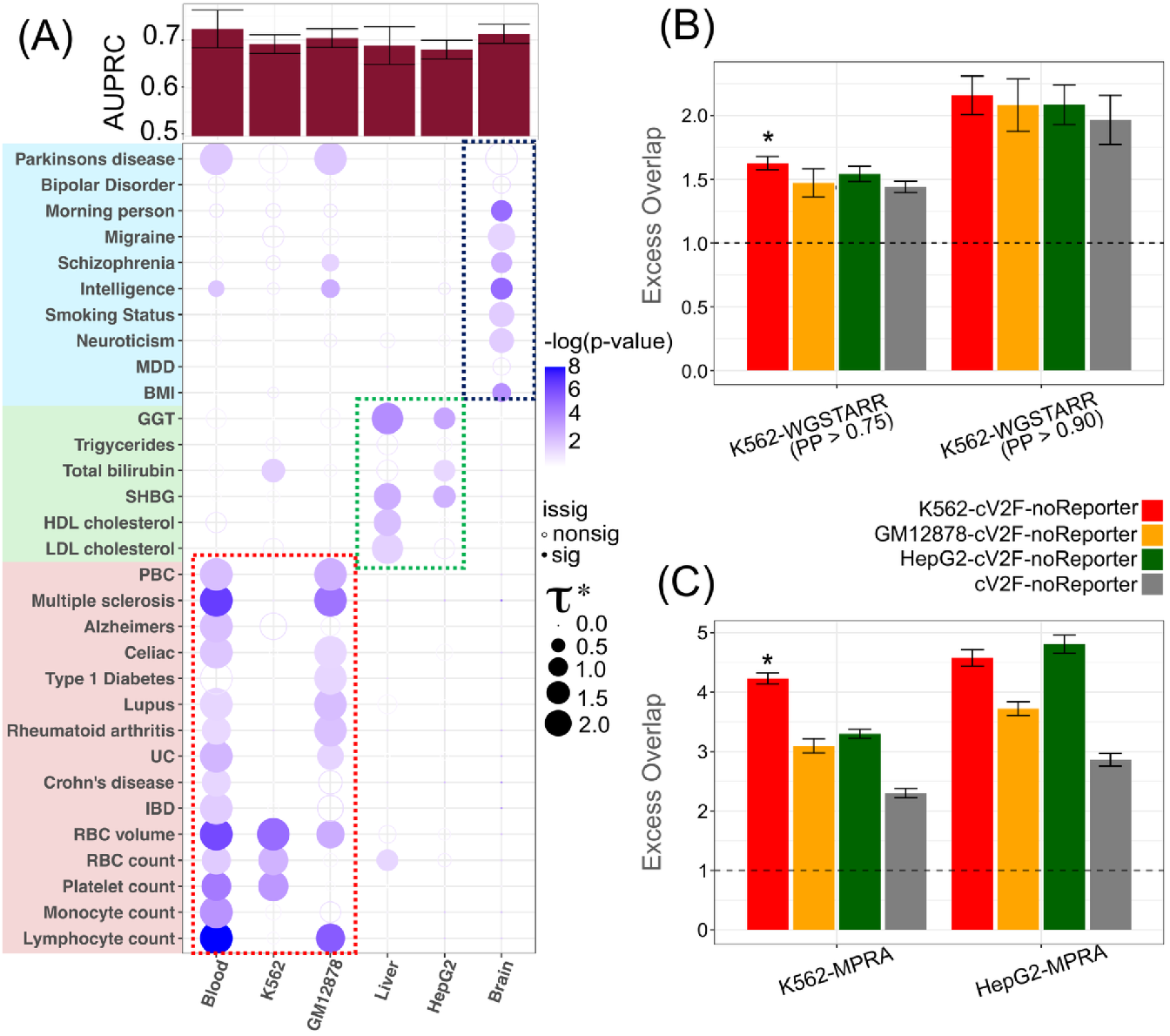
Properties of cell-line and tissue-specific cV2F scores. (A) (Top panel) Area under the Precision Recall curve (AUPRC) of cell-line and tissue-specific cV2F gradient boosting models based on held-out GWAS fine-mapping data. (Bottom panel) S-LDSC standardized effect sizes (τ*) of cell-line and tissue-specific cV2F scores for a set of related diseases and traits. Results are conditional on 97 baseline-LD v2.2 + 1 binarized primary cV2F annotations. Magnitude (***τ***\*, dot size) and significance (−log10(*P*), dot color) are reported for disease signal for 31 blood, liver and brain-related traits. (B) Excess overlap of binarized cell-line specific binarized cV2F-noReporter for three cell lines - K562, GM12878, and HepG2 (colored), and primary cV2F-noReporter (gray) against Whole Genome STARR-seq positives versus tested variants in K562. For the STARR-seq assay, two thresholds on posterior probability of allelic effect (0.75 and 0.9) from ref^116^ were considered. (C) Excess overlap of binarized cell-line specific binarized cV2F-noReporter (model trained without MPRA features) variants for three cell lines - K562, GM12878, and HepG2 (colored), and primary cV2F-noReporter (gray) against MPRA positives versus tested variants in related cell lines (K562 and HepG2). Error bars denote 95% confidence intervals. Numerical results are reported in **Supplementary Table S7**.

We assessed the (binarized) tissue/cell line-specific cV2F scores using the S-LDSC disease heritability framework^28,47^. To assess whether the tissue/cell line-specific cV2F scores contain unique information, we conditioned on the (binarized) primary cV2F score as well as 97 baseline-LD (v2.2) model annotations^27^ (baseline-LD-cV2F model). We observed substantial heritability enrichments across 31 matched complex traits related to blood, liver and brain (average 15.6x, s.e. 2.3); these enrichments were comparable to the primary cV2F score on the same traits (average 17.6x s.e. 0.40)(**Supplementary Figure S8B, Supplementary Table S7**). Importantly, we observed large and Bonferroni-significant ***τ***\* values for diseases/traits corresponding to these tissues/cell lines (**Figure 4A bottom panel, Supplementary Table S7)**; as noted above, annotations with ***τ***\* value >0.5 are generally considered to be highly important for disease^64,83^. For example, Liver-cV2F attained ***τ***\* = 1.81 for LDL and ***τ***\* = 1.35 for HDL, and and GM12878-cV2F attained ***τ***\* = 1.57 for lymphocyte count. We observed large and highly significant ***τ***\* values when meta-analyzing ***τ***\* values across traits related to each tissue/cell line, e.g. ***τ***\* = 1.09 (p = 1.7e-09) for Blood-cV2F across 15 blood-related traits (**Supplementary Figure S8C**). We determined that using only tissue/cell line-related traits for training led to lower ***τ***\*, likely due to insufficient amount of training data (**Supplementary Figure S9**).

We assessed the performance of the (binarized) tissue/cell line-specific cV2F scores with respect to variants implicated by the MPRA^12^ and WG-STARR-seq assays; as above, we removed MPRA features from cV2F in this analysis (cV2F-noReporter). For variants implicated by WG-STARR-seq available in K562, K562-cV2F-noReporter outperformed other tissue/cell line-specific cV2F scores as well as the primary cV2F score (**Figure 4B** and **Supplementary Table S7**). For variants implicated by MPRA in K562 and HepG2, the corresponding cell line-specific cV2F scores generally outperformed other tissue/cell line-specific cV2F scores as well as the primary cV2F score (**Figure 4C** and **Supplementary Table S7**). These results indicate that tissue/cell line-specific cV2F is effective in capturing tissue/cell line-specific information (despite the fact that MPRA experiments are known to incompletely mimic the native cell-line specific chromatin context^84,85^).

We used the binarized Blood-cV2F and Liver-cV2F scores to perform functionally informed fine-mapping of variants associated with 19 blood-related traits and 7 liver-related traits^86^ (subselected from the 110 UK Biobank traits), by reweighting posterior probabilities as above. We observed a 7.1% increase for Blood-cV2F and 9.1% increase for Liver-cV2F in the number of confidently fine-mapped (PIP > 0.95) variants; however, this increase was lower than the improvement for the primary cV2F score for the same traits (11.3% and 13.5% respectively) (**Supplementary Tables S6, S8, S9**), consistent with the smaller number of features by Blood-cV2F (47) and Liver-cV2F (30) compared to the primary cV2F score (339). When restricting to blood-related and liver-related traits, Blood-cV2F-informed and Liver cV2F-informed fine-mapping identified 843 and 502 confidently fine-mapped variants (PIP>0.95) (**Supplementary Tables S8, S9**). We conclude that tissue and cell-line specific cV2F scores show unique disease heritability information conditional on the primary cV2F score and show specific enrichment in functional characterization experiments in matched biosamples.

### Leveraging cV2F to pinpoint causal GWAS variants with high confidence

We highlight 4 GWAS loci for which functionally informed fine-mapping using cV2F pinpoints a causal variant with high confidence (PIP > 0.95). First, at the *PLAG1-CHCHD7* locus for birth weight, the PIP for *rs72656010* (GWAS p-value = 2e-26) improves from 0.80 to 0.97 when incorporating the cV2F score; this variant is implicated by ABC, cCRE, Enformer and Zoonomia (**Methods, Figure 5A**). This variant is linked by the cS2G method^79^ to the *PLAG1* and *CHCHD7* genes (with similar scores). Of these two genes, *PLAG1* has been previously implicated for growth-related traits in animal studies, including knockout studies in mice^87,88^; this implicates *PLAG1* as a likely causal gene at this locus.

**Figure 5:**
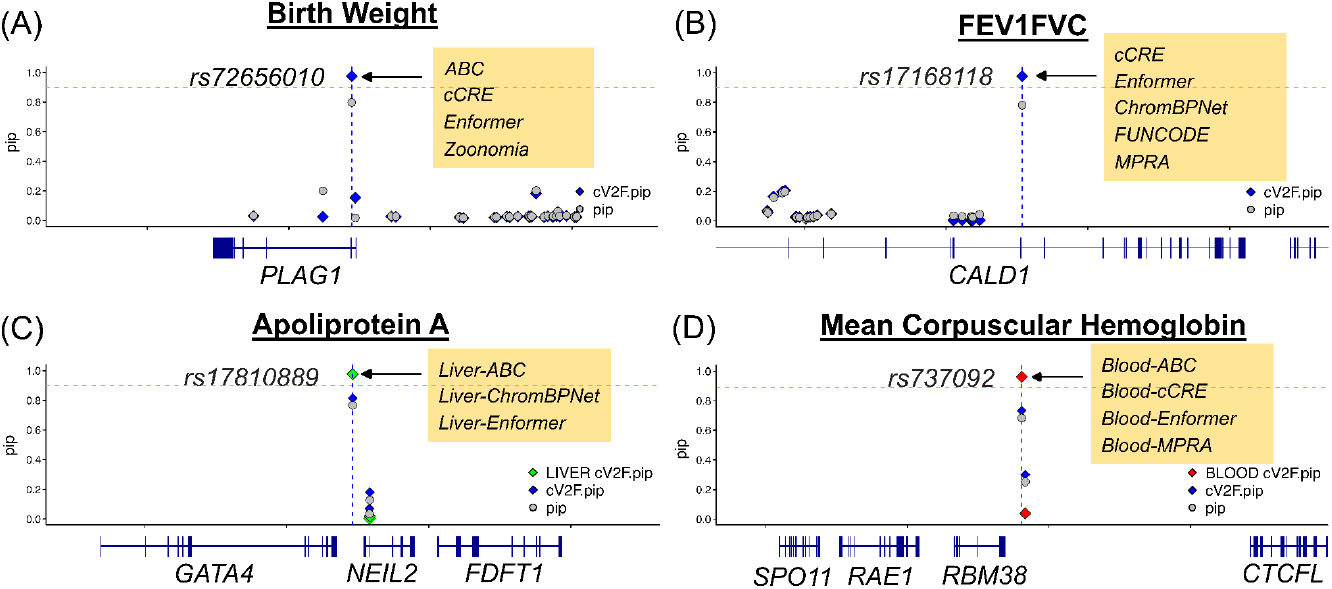
Examples of GWAS variants implicated by cV2F fine-mapping. We illustrate 4 confidently fine-mapped variants implicated by fine-mapping informed by primary cV2F or tissue-specific cV2F scores. (A) *rs72656010* is implicated for Birth weight (cV2F.PIP=0.97) and (B) *rs17168118* is implicated for the lung capacity trait FEV1/FVC (cV2F.PIP=0.98) by primary cV2F fine-mapping but not standard fine-mapping, (C) *rs17810889* is implicated for Apoliprotein A (Liver.cV2F.PIP=0.98) by Liver-cV2F fine-mapping but not by primary cV2F fine-mapping or standard fine-mapping, (D) *rs737092* is implicated for Mean corpuscular hemoglobin (Blood.cV2F.PIP=0.96) by Blood-cV2F fine-mapping but not by primary cV2F fine-mapping or standard fine-mapping. Selected features for which the implicated variant scores in the top 5% are reported in boxes.

Second, at the *CALD1* locus for lung capacity trait FEV1/FVC, denoting the ratio of FEV1 (Forced Expiratory Volume) and FVC (Forced Volume Capacity), the PIP for intronic variant *rs17168118* (GWAS p-value = 7e-10) improves from 0.78 to 0.98 when incorporating the primary cV2F score; this variant is implicated by cCRE, Enformer, ChromBPNet, FUNCODE and MPRA features (**Figure 5B**). This variant is strongly linked to the *CALD1* gene (cS2G=1)^79^ (**Supplementary Table S6**). The *CALD1* gene is known to regulate smooth muscle and non-muscle contraction and has been previously implicated as a dysregulated hub gene in TGF-β1-induced pulmonary fibrosis model^89^.

Third, at the *GATA4/NEIL2* locus for Apolipoprotein A, the PIP for *rs17810889* (GWAS p-value = 2e-15) improves from 0.78 to 0.98 when incorporating the Liver-cV2F score (vs. only 0.81 when incorporating the primary cV2F score); this variant is implicated by Liver-ABC, Liver-ChromBPNet and Liver-Enformer features (**Figure 5C**). This variant is strongly linked to the *NEIL2* gene (cS2G=1)^79^. *NEIL2* encodes DNA glycosylase enzymes, and recent studies implicate this gene in APOBEC3-mediated mutagenesis^90,91^.

Fourth, at the *RBM38* locus for mean corpuscular hemoglobin (MCH), the PIP for *rs737092* (GWAS p-value = 8-e134) improves from 0.72 to 0.96 when incorporating the Blood-cV2F score (vs. only 0.72 when incorporating the primary cV2F score); this variant is implicated by the MPRA data-driven deep learning feature (K562) and blood-specific ABC, cCRE and Enformer features (**Figure 5D**). Among blood cell lines, K562-cV2F-informed fine-mapping showed higher confidence of function at this variant compared to GM12878 (PIP=0.98 vs. 0.76). This variant has been strongly linked to the *RBM38* gene by the cS2G method (cS2G=0.96)^79^. Previous MPRA showed allele-specific activity for this variant in K562+GATA1, and targeted deletion of this variant in K562 showed strongest effect on the expression of the *RBM38* gene^92^. *RBM38* is an erythroid-specific RNA binding protein that plays a critical role in terminal erythroid differentiation, consistent with playing a role in hemoglobin regulation^93,94^.

We conclude that incorporating the primary and tissue/cell line-specific cV2F scores into fine-mapping identifies and functionally characterizes causal GWAS variants with high confidence, recapitulating known biology.

## Discussion

We have proposed a consensus variant-to-function (cV2F) score that prioritizes variants for disease-related function by integrating GWAS data with a broad range of variant-level and element-level functional information from experimental assays and sequence-based deep learning models. We also propose tissue/cell line-specific cV2F scores that leverage tissue/cell line-specific functional information. We determined that the cV2F score contains unique information about disease, outperforms other pathogenicity scores^25,26,58–60,63,65–68,70–72,74–77^ across a range of metrics, and improves statistical fine-mapping. We note that the cV2F score expands upon an unpublished preprint^95^ which contained related ideas but was focused on integrating information from deep learning models; here, we integrate a richer array of functional information, consider a broader set of validation metrics, construct tissue/cell line-specific scores, and investigate incorporation into fine-mapping.

Our work has several downstream implications. First, our primary and tissue/cell line-specific cV2F scores can improve statistical fine-mapping in GWAS data beyond the UK Biobank data that we have analyzed here; in particular, we anticipate considerable utility for multi-ancestry fine-mapping^54,96–98^, as functional architectures of disease are generally consistent across ancestries (**Supplementary Figure 3C** and ref.^54–56^). Second, our cV2F scores and the fine-mapped disease variants that they implicate can inform future functional follow-up experiments such as MPRA^15,85^, CRISPRi^14,99^ and CRISPR base editing^100,101^ experiments. Third, our cV2F scores can also be used to improve functionally informed polygenic risk prediction within and across ancestries^102,103^; as described in a companion manuscript^104^.

Our work has several limitations, representing important directions for future research. First, we restricted our focus to specific element-level and variant-level features from ENCODE4; there is potential to extend the model by including more extensive feature sets capturing silencer elements^105,106^, DNA methylation^107^, post-transcriptional modifications^108^ or molecular QTL (beyond eQTL, e.g. chromatin QTL or protein QTL^109,110^). Additional features could capture enhancers and regulatory elements capturing the dynamic nature of cell systems, such as inflammation, stimulation, development systems and *trans* effects of perturbations^36,37,111–113^. Second, our tissue/cell line-specific cV2F scores currently span a limited set of tissues and cell lines motivating the generation of functional data in other tissues, cell lines, and fine-grained cell types (e.g. in single-cell data^114^). Third, we use fine-mapped variants for a broad set of UK Biobank diseases/traits as training data for both primary and tissue/cell line-specific cV2F scores (due to the limited number of fine-mapped variants for a single disease/trait), precluding disease-specific inference. Incorporating fine-mapped variants from a limited set of genetically correlated diseases/traits^115^ could be a route towards disease-specific inference. Fourth, we primarily focused on binarized cV2F scores (due to the non-linear relationship between excess overlap of fine-mapped GWAS variants and cV2F probabilistic score; **Figure 2C**). Binarizing probabilistic scores loses information, such that alternative approaches could potentially be more powerful. Fifth, we have primarily trained and validated the cV2F model using data from European-ancestry cohorts. However, analyses of other ancestries (**Supplementary Figure S3**) suggest that our cV2F model can be used to score variants in non-European populations. Sixth, we performed functionally informed fine-mapping incorporating primary or tissue/cell line-specific cV2F scores using a post-hoc approach (**Figure 3E**), but incorporating cV2F scores and other annotations into a formal functionally informed fine-mapping framework remains as a future direction. We have shown that the post-hoc approach using cV2F slightly outperforms PolyFun + SuSiE^6^ using the baseline-LD model (14.3% improvement vs. 10.7% improvement compared to non-functionally informed fine-mapping, **Supplementary Table S6)**, but incorporating cV2F into PolyFun + SuSiE may confer a greater benefit, particularly as our heritability enrichment analyses show that cV2F scores are uniquely informative for disease (**Figure 3A** bottom panel, **Figure 4A** bottom panel). Despite all these limitations, our primary and tissue/cell line-specific cV2F scores are a promising approach to functionally prioritize genetic variants for disease.

## Methods

### Genomic annotations in the cV2F model

We define a functional annotation as an assignment of a numeric value to each SNP with minor allele count >=5 in the European cohort reference panel from the 1000 Genomes Project. Annotations can be either binary or continuous-valued. In the primary (tissue-agnostic) cV2F model, we considered 339 features encompassing a broad range of functional assays and models. These 339 features include 186 variant-level annotations, 52 genomic element-level features, 84 previously established disease-related features from the baseline-LD model^28,47^, and 17 additional functional and sequence-based conservation-related features from ENCODE4 and Zoonomia^29,30^. For the cell-line/tissue-specific cV2F model, we considered subsets of features relevant to the specific cell line or tissue of interest. For Blood-cV2F, we retained 47 features aggregated over blood biosamples, as well as the related K562 and GM12878 biosamples. For Liver-cV2F, we retained 30 features aggregated over liver biosamples, as well as the related HepG2 biosample. For Brain-cV2F, we retained 19 features aggregated over brain biosamples. For K562-cV2F, GM12878-cV2F and HepG2-cV2F, we retained 16, 12, and 15 features corresponding to K562, GM12878, and HepG2 biosamples respectively.

### Variant-level functional annotations in the cV2F model

The 186 variant-level functional annotations were broadly derived from 9 different data sources, representing biochemical assays, sequence-based models and quantitative trait loci (QTLs). The processing of these annotations is detailed as follows.

#### Transcription Factor Allele-Specific Binding

We curated 15 Transcription Factor (TF) Allele-specific binding event features from ADASTRA Bill Cipher version (v5.1.3, 2022.05.19)^39^ corresponding to 2,180 TF and cell-type pairs. For 15 cell lines and tissues (**Supplementary Table S1**), we created a feature annotation defined by variants with allele-wise logit-aggregated and FDR-corrected P-values <10% significance for imbalance at the reference and alternate alleles for at least one TF in that cell line/tissue.

#### DNase Allelic Imbalance

We curated 28 DNase allelic imbalance event features from ENCODE allelic imbalance results (version 5) across 3,712 biosamples (J. Vierstra unpublished data, **Data Availability**) (**Supplementary Table S1)**. For each of 15 cell lines and tissues, we considered two variant feature annotations given by variants that are (i) normally significant minimum p-value < 0.05 or (ii) minimum FDR<10% significant between reference and alternate alleles. Additionally, we used the “all.aggregated.bed” results to call nominal and FDR<10% allelically imbalance variants across all biosamples (see Data Availability). Features corresponding to biosamples for which the number of variants implicated at nominal or FDR significance was <100 were dropped. Variants that were not tested were assigned a mean imputed value based on the number of significant variants.

#### Reporter assays

We considered 6 variant features derived from two reporter assays - a Massively Parallel Reporter Assay (MPRA) experiment and a Whole-Genome STARR-seq assay (**Supplementary Table S1)**. The MPRA data targeted 180, 781 GTEx fine-mapped eQTLs across 49 tissues in 4 cell lines (HCT116, HepG2, K562, SK-N-SH)^13^. An expression smodulating variant (emVar) was defined as variants residing in active elements with any magnitude of allelic effect, an activity cut-off given by absolute value of log2-fold-change >=1 for either of two alleles, an activity FDR given by log-P-adjusted-bayesFactor >=2 for either of the two alleles, the control read depth > 50 at both alleles, and an FDR < 10% for a non-zero allelic effect. The Whole-Genome STARR-seq assay in K562 cell line targeted natural genetic variation in African population subgroups (T. Reddy, unpublished data, **Data Availability**); we considered two thresholds (0.75 and 0.90) on the posterior probability of allelic effect from the BIRD method^116^. Variants that were not tested were assigned a mean imputed value based on the number of significant variants.

#### Sequence-based computational models

We considered 108 annotations corresponding to deep learning models trained on DNA sequence data targeted at predicting chromatin activity, histone marks and reporter assay enhancer activity (**Supplementary Table S1**). We consider two sequence-based deep learning frameworks trained on ENCODE biochemical assay tracks - Enformer, a multi-tasking model across many cell types and features, ChromBPNet^40,41^, a cell-type resolution Tn5 bias-corrected convolutional model, and a convolutional neural net deep learning model trained on MPRA enhancer activity by R. Tewhey Lab. We compiled predicted allelic activity of each of 9,991,229 variants in consideration for each biosample and track using these models. The Enformer model annotations were generated for all variants for each of 5,313 features. These features were then converted to a Z-score scale by standardizing across all variants, and features corresponding to similar assay type and cell type were combined by taking the maximum standardized score. This resulted in 60 Enformer annotations corresponding to 15 diverse cell types and feature categories (all, CHIP, DNase, H3K4me3, H3K4me1, H3K27ac). Certain assay and cell-type pairs were dropped if the size of the resulting annotation is less than 0.001%.

For the ChromBPNet model (A. Kundaje, unpublished data, **Data Availability**), predicted allelic effects were generated for all variants for 101 DNase biosamples, and three metrics were computed for each biosample - (i) average cross-product of absolute log-fold change (*abs*.*logFC*), (ii) Jensen Shannon Divergence (*jsd*), and (iii) their cross-product (*abs*.*logFC x jsd*) with maximum percentile mean activity. Each metric was filtered based on the p-value of the cross product being less than 0.05. For each of these metrics, we aggregate features by taking the maximum across biosamples related to 15 focal cell types and tissues; this resulted in a total of 36 features. For the MPRA-based deep learning model, we considered 12 features corresponding to the average of allelic skew predictions between reference and alternate alleles, and just binarized versions of these scores based on crossing the p-value<0.05 significance cut-off for reference and alternate alleles in 3 cell lines (K562, GM12878 and SK-N-SH).

For the MPRA deep learning model (R. Tewhey, unpublished data, **Data Availability**), 10 cross validated models were generated for eleven autosome pairs such that every chromosome has ten models in which it has been held out from training. MPRA model predictions were generated for variants in a chromosome specific manner and the average of the ten model outputs was taken to give a final activity score. Variant effect scores represent the difference in activity between the alternate and reference allele sequences. To better capture variant effects, 18 ‘windowed’ predictions were generated for each variant wherein the variant position within the tested sequence shifted by 10 bp in each prediction. These 18 predictions were averaged to give a variant effect score. As the model predicts for MPRA activity in three cell lines (K562, HepG2, SKNSH), the variant effect score of greatest absolute effect was used as the final score.

#### QTL features

We included 29 quantitative trait loci (QTL) features from GTEx and the ENCODE AFGR project (**Supplementary Table S1)**. We considered 20 GTEx fine-mapping variant features, binarized at two posterior inclusion probability (PIP) thresholds 0.10 and 0.50, for at least one cis-eGene across 9 human tissues^38^ (we also consider 2 features corresponding to the union of these fine-mapped eQTLs pooled across all tissues). We considered these thresholds as they have been previously used in eQTL benchmarking efforts^34,38^. We also considered 9 annotations from the African Functional Genomics Resource (AFGR) project that included nominal (p-value < 0.05) and FDR < 5% significant (using both Bonferroni and Benjamini Hochberg methods significant eQTL and caQTL annotations in the Lymphoblastoid Cell Line (LCL), as well as variants with moderate to high PIP by considering binarizing at three thresholds (PIP> 0.10, 0.50 and 0.75) from single causal variant fine-mapping performed on the eQTL data.

### Other annotations in the cV2F model

Outside of the variant-level annotations, the cV2F model includes 52 element-level annotations comprising of 40 cCRE (v4) element features^10^, both primary and tissue-specific, and capturing 3 broad categories of signatures (PLS: Promoter-like, pELS: Proximal Enhancer-like and dELS: Distal enhancer-like) and 12 Activity-By-Contact (ABC) enhancer element features across 9 different tissues (blood, brain, kidney, liver, heart, lung, intestine, fat, skin), assessed as critical for some complex diseases and traits^24^, and 5 ENCODE cell-lines (K562, GM12878, HepG2, SKNSH and A549) (**Supplementary Table S1)**. For cCRE elements, we considered the union of related biosample-level annotations across 1678 biosamples into relevant tissue and cell-types. For ABC elements, we considered the union of biosample-level element-gene links with ABC score > 0.015 across 352 biosamples into relevant tissues and cell types. We used the ABC method to implicate the elements over the recently proposed ENCODE-rE2G^34^ approach despite superior performance of the latter, since ENCODE-rE2G uses a logistic regression-informed weighting of element-gene links from all chromosomes, which may lead to possible overfitting in the cV2F model. Cell lines and tissues for which <1% variants were annotated by the element-function strategy were dropped from the feature set. Additionally, the cV2F model includes 84 annotations from the baseline-LD (2.2) annotations that span coding, conserved, and broader regulatory annotations^27^ (we do not consider MAF bin and LD related annotations as we correct for these effects when training the cV2F model) (see **Supplementary Table S1**).

In the cV2F model, we incorporated 19 additional conservation annotations, separate from the ones included in baseline-LD, comprising of 17 FUNCODE functional conservation^44^ (**Data Availability)** scores and 2 Zoonomia sequence-based conservation across species^29,30^. The FUNCODE (FUNctional COnservation of DNA Elements) framework was developed to quantify functional conservation between human and mouse regulatory elements using chromatin accessibility and histone modification data from the ENCODE4 project^44^. This approach leverages a comprehensive compendium of functional genomic data from both species to assess two distinct aspects of conservation: tissue/cell-type-specific activities (CO-V) and constitutive baseline activities (CO-B). CO-V is designed to capture the conservation of regulatory elements that exhibit dynamic, context-specific activity patterns across various tissues or cell types. To achieve this, FUNCODE employs an unsupervised sample matching technique to identify corresponding tissue/cell types across species. This method enables more efficient use of existing datasets without the need for manual annotation. CO-V is computed using weighted Spearman’s correlation statistics of activities across matched samples between species. In contrast, CO-B aims to identify regulatory elements that maintain consistently high activity levels across most contexts in both species, such as promoters or enhancers of housekeeping genes. CO-B is calculated using a statistic inspired by the h-index, which does not require sample matching, making it well-suited for capturing elements with high baseline activity.FUNCODE was applied to 2,595 uniformly processed ENCODE experiments, generating genome-wide conservation scores for 2.24 million sequence-aligned pairs. The resulting FUNCODE-conserved elements showed significant enrichment in disease-associated GWAS variants. Notably, although defined based on human-mouse conservation, the score demonstrated the ability to identify functionally conserved elements between more distantly related species, such as human and zebrafish. Unlike traditional sequence-based conservation scores, FUNCODE captures the conservation of regulatory element function and activity across species.

### Training data for the cV2F model

We use for training data a set of positive variants that are confidently fine-mapped variants (PIP > 0.90) across 94 traits in UKBiobank as used in ref^34,52^. We considered as negatives 1,227,661 variants that have very low fine-mapping posterior probability (PIP < 0.01) for all 94 UKBiobank traits. In order to ensure better matching between the positive and negative set of variants, we restrict our negative set of variants to the ones that are in the same LD block^117^ as a positive variant and up to 5 closest variants in terms oft Minor Allele Frequency (MAF) as the positive variant. We assessed the relative performance of different features based on the Area Under the Precision Recall Curve (AUPRC) of the predicted probabilities of fine-mapping against the positive and negative set of variants from held-out chromosomes. We considered other PIP thresholds for the set of positive variants (PIP=0.1, 0.25, 0.5, 0.75), however the overall AUPRC for these choices was lower (**Supplementary Table S2**); this highlights that higher confidence of fine-mapped signals results in an improved cV2F prediction model. As secondary analysis, we also considered training using 5,269 confidently fine-mapped variants (PIP> 0.9) from 931 traits in Million Veteran Program (MVP) project against 90,532 negative variants (PIP < 0.01) that were then LD and MAF matched to the positive variant. We used the same training data for cell-line and tissue-specific versions of cV2F, but also considered for secondary analyses, training data regimes restricted to traits and diseases matched to the related cell lines and tissues.

### Generating the cV2F score

We used the Extreme gradient boosting (XGBoost) method implemented in the XGBoost software^32,118^ with the following model parameters: the number of estimators (25, 40, 50), depth of the tree (10, 15, 25), learning rate (0.05), gamma (minimum loss reduction required before additional partitioning on a leaf node; 10), minimum child weight (6, 8, 10), and subsample (0.6, 0.8, 1); we optimized parameters by tuning hyper-parameters (a randomized search) with five-fold cross-validation. Two important parameters to avoid over-fitting are gamma and learning rate; we chose these values consistent with previous studies^77^, as in our previous work on AnnotBoost^24^ and DeepBoost^95^ frameworks.

The gradient boosting predictor is based on T additive estimatorsand it minimizes the loss objective function *L_t_* at iteration *t*.

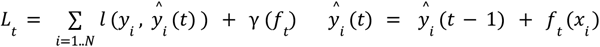

Where *f*_*t*_ is an independent tree structure and γ (*f*_*t*_) is the complexity parameter. The final prediction from the gradient boosting model therefore is given by

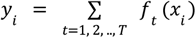

In order to avoid winner’s curse and overfitting, we use fine-mapped SNPs on odd (respectively even) chromosomes as training data to make predictions for even (respectively odd) chromosomes, as in our previous work on AnnotBoost^24^; thus, boosted annotations on a given chromosome are not informed by fine-mapped SNPs on that chromosome. We report the average AUPRC (Area under Precision Recall Curve) of odd and even chromosome classifiers. The boosted annotations produced as output of the classifier are probabilistic in nature because of the logistic loss. We have performed independent training and prediction generation for a model with all features (variant-level, element-level, baseline-LD), as well as different subsets of these features. The importance of each feature is assessed using Shapley scores^50^ and marginal AUPRC of each feature in the model.

### Excess overlap and recall metrics for the cV2F score

We reported excess overlap of variants annotated by binarized V2F method against validation set of variants from experimental assays, GTEx fine-mapping and GWAS fine-mapping data. For a given V2F method, we define *V*2*F*(*annot*) as a binary set of variants annotated by the V2F method in terms of passing a score threshold (0.75 for cV2F) and *V*2*F*(*total*) represents the total number of variants scored by it, which totals to 9,991,229. On the other hand, for a given validation data (*valid*), we define *valid*(*sig*) as the number of significant variants in the validation data set, passing specific significance cut-offs and log-fold-change or other related thresholds, as prescribed for each method. Additionally, *valid*(*tested*) represents the number of variants actually tested or passing the quality check (QC) for the given assay. Then we define the excess overlap metric as follows

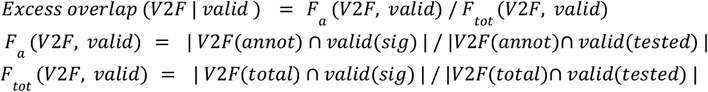

Similarly, we define the recall metric as follows

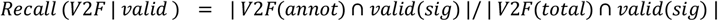

The standard error of the excess overlap and recall metrics are calculated by Jack-knifing over LD blocks. We considered a variety of experimental assays for these metrics - CRISPRi data in K562, MPRA data in 5 cell lines, WG-STARR-seq data in K562. We have also considered this metric against a validation set of GWAS fine-mapped variants and GTEx fine-mapped eQTLs. The variant universe of each validation dataset was restricted to the set of the set of variants scored by the V2F scores, which equals the 9,991,229 variants (hg38) with minor allele count >=5 in 1000 Genomes Europeans. For the CRISPR data, we considered 1,185 and 22,124 variants located within 1KB of significant and tested peak-gene links from 3 CRISPRi experimental assays in K562 respectively^35–37^. For the MPRA data, we considered 16,175 and 136,173 variants significant and tested variants for enhancer activity across 5 cell lines, where the tested variants were drawn from GWAS fine-mapped variants in Biobank Japan^56^ and UKBiobank^52^; for the cell-type specific version, we restricted the significant variants to the focal cell line. For the WG-STARR data, we considered 5,873 (1,752) and 204,802 variants significant (BIRD^116^ posterior probability > 0.75 (0.90)) and tested variants for enhancer activity in K562.

We compared cV2F against Regulomedb v2.2 scores^58^; for the primary comparison, we used variants with Regulomedb probability score > 0.90; but we additionally also compared variants in different categories of Regulomedb score (**Figure 3A, Supplementary Figure S4**). We compared against MACIE scores^60^; for the primary comparison, we considered MACIE-anyclass score > 0.99 but for secondary comparison we also tested against MACIE-conserved and MACIE-regulatory scores (**Figure 3A, Supplementary Figure S4**). For the pathogenicity scores comparison, we downloaded primary and secondary pathogenicity scores and their boosted versions from ref^24^, where thresholds for each score were determined based on optimal disease heritability enrichments.

### Stratified LD score Regression

Stratified LD score regression (S-LDSC) is a method that assesses the contribution of a genomic annotation to disease and complex trait heritability^28,57^. S-LDSC assumes that the per-SNP heritability or variance of effect size (of standardized genotype on trait) of each SNP is equal to a linear contribution of each annotation. 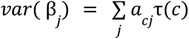, where *a*_*cj*_ is the value of annotation c for SNP j, and τ(*c*) is the contribution of annotation c to per-SNP heritability conditioned on other annotations. S-LDSC estimates the τ(*c*) for each annotation using the following equation:

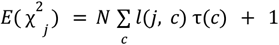

where 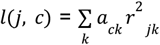 is the stratified LD score of SNP j with respect to annotation c and *r*_*jk*_ is the genotypic correlation between SNPs j and k computed using data from 1000 Genomes Project (see URLs); N is the GWAS sample size. We assess the informativeness of an annotation c using two metrics. The first metric is enrichment (E), defined as follows (for binary and probabilistic annotations only):

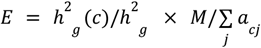

*wh*ere 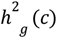 is the heritability explained by the SNPs in annotation c, weighted by the annotation values. The second metric is standardized effect size (τ*) defined as follows:

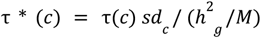

where *sd*_*c*_ is the standard error of annotation c, *h*^*2*^ _*g*_ is the total SNP heritability and M is the total number of SNPs on which this heritability is computed (equal to 5,961,159 in our analyses). τ * (*c*) represents the proportionate change in per-SNP heritability associated to a 1 standard deviation increase in the value of the annotation.

### Functional fine-mapping using the cV2F score

We perform a functional fine-mapping of UKBiobank GWAS associations using the cV2F score following a similar approach as in Expression Modifier Score (EMS)^38^. The SuSIE fine-mapping model^5^ assumes a causal configuration vector 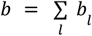 where both *b* and *b*_*l*_ are vectors of length m, the number of variants in the locus to be fine-mapped, and *b*_*l*_ has one entry equals to 1 and all other entries as 0. SuSIE defines a set of vectors (of length *m*) α_*1*_, α_*2*_, …., α_*L*_ with α_*l*_ (*v*) = *Pr* (|*b*_l_ (*v*)| > 0 | *X*) is the posterior probability given the data *X*. Credible sets are computed for each *l* from α_*l*_ and credible sets that are not pure, meaning that contain a pair of variants with absolute correlation < 0.5, are pruned out. If α_*l*_ corresponds to a pure credible set, we re-weight each element of α_*l*_ with the binarized primary or tissue-specific score. Let *c*_*1*_, *c*_*2*_, …., *c*_*m*_ be the cV2F scores for *m* variants, then we generate updated posterior probabilities δ_*l*_ as

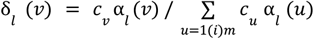

We use these updated cV2F-informed posterior probability δ_*1*_, δ_*2*_, …., δ_*L*_ to compute updated credible sets and posterior inclusion probabilities (cV2F.PIP). We compare these cV2F.PIP values with the standard PIP values obtained from the non-functionally-informed posterior probabilities α_*l*_ (*v*). For the tissue-specific approach, we replaced the cV2F score with tissue or cell-line cV2F score. For the analysis in the paper, we focused on two tissues - Blood and Liver, since for these two tissues, several related biomarker traits with well-powered fine-mapping were available.

### Functional features implicated at a variant

For a selected variant by the cV2F functional fine-mapping as highlighted in the locus plots in Figure 5, we nominate features as “implicated” for the variant in the following way. For a binary 0/1 feature, we implicate it for the variant if the variant is annotated as 1 by that feature. For a continuous feature, we implicate it for a variant if the variant is in the top 1% of variants with the highest value of the quantitative score for the focal feature.

## Data Availability

cV2F scores (**Supplementary Table S3**): https://mskcc.box.com/shared/static/hsrogtr3fddtmd53hyy5ph7dlp20eq72.txt UKBiobank fine-mapping data: https://www.finucanelab.org/data and gs://finucane-requester-pays/ukbb-finemapping/MVP fine-mapping data: Verma et al.^53^

https://drive.google.com/file/d/1XSyAgpCXQTFXHyHnR6gChXmzB6p9nEzg/view

https://drive.google.com/drive/folders/1SKm5caoFziLBu5VST2zt0TevbMLNRHqm

GTEx eQTL fine-mapping data: https://www.finucanelab.org/data AFGR QTL data: https://github.com/smontgomlab/AFGR Activity-By-Contact (ABC) maps:

https://www.encodeproject.org/search/?type=Annotation&annotation_type=element+gene+regulatory+interaction+predictions&status=released&software_used.software.name=distal-regulation-encode_re2g; see “derived from” field of https://www.encodeproject.org/metadata/?type=Annotation&annotation_type=element+gene+regulatory+interaction+predictions&status=released&software_used.software.name=distal-regulation-encode_re2g to obtain ABC predictions files. MPRA data: R. Tewhey lab: ENCODE portal (see accession IDs from Supplementary Tables S25, S26 in ref^13^).

FUNCODE scores: ENCODE portal

(https://www.encodeproject.org/search/?type=Annotation&annotation_type=cross-species+functional+conservation). FUNCODE score track hubs for the UCSC genome browser are available on GitHub (https://github.com/wefang/funcode). FUNCODE scores for regulatory elements are accessible by query through the web application (https://jhubiostatistics.shinyapps.io/FUNCODE).

ChromBPNet scores (Kundaje lab), Whole-genome STARR-seq data (Reddy lab), MPRA allelic skew effects and deep learning model annotations (Tewhey lab), DNase Allelic imbalance calls (Viertsra lab),and cCRE (v4) maps (Moore, Weng labs) informing cV2F features are ENCODE (Phase 4) datasets that are currently available via request to the authors and will be made publicly available on the ENCODE portal (https://www.encodeproject.org/) prior to the publication of this manuscript.

## Code Availability

Code links to replicate cV2F scores, metrics and finemapping are available on GitHub (https://github.com/Deylab999MSKCC/cv2f).

## Supplementary Figures

**Supplementary Figure S1:**
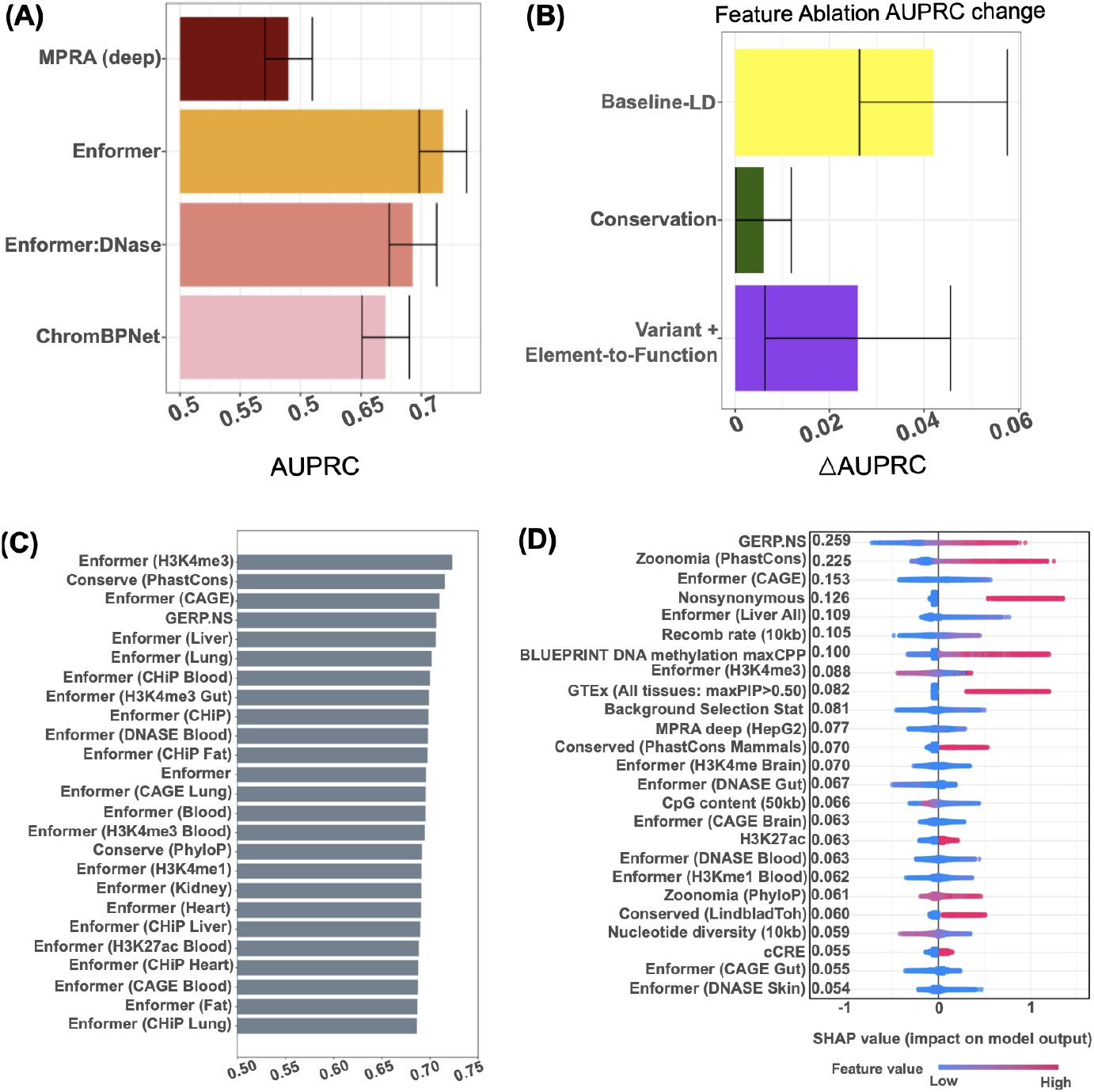
(A) The AUPRC of the GWAS fine-mapping gradient boosting training model corresponding to deep learning predicted allelic effect functional annotations from MPRA-based deep learning model, Enformer, Enformer restricted to DNase features and ChromBPNet features. (B) Change in Area Under the Precision Recall Curve (AUPRC) of the cV2F GWAS fine-mapping based gradient boosting training model upon ablation of three broad categories of functional annotations - baseline-LD, conservation annotations (Zoonomia and FUNCODE) and ENCODE element and variant-level functional annotations. (C) The AUPRC of the gradient boosting training model for individual functional features; results are reported for the top 25 features. See Supplementary Table 2 for results from the full set of 339 features. (D) The Shapley values of the top 25 predictive features based on the cV2F gradient boosting training model. Error bars denote 95% confidence intervals. Numerical results are reported in **Supplementary Table S2**.

**Supplementary Figure S2:**
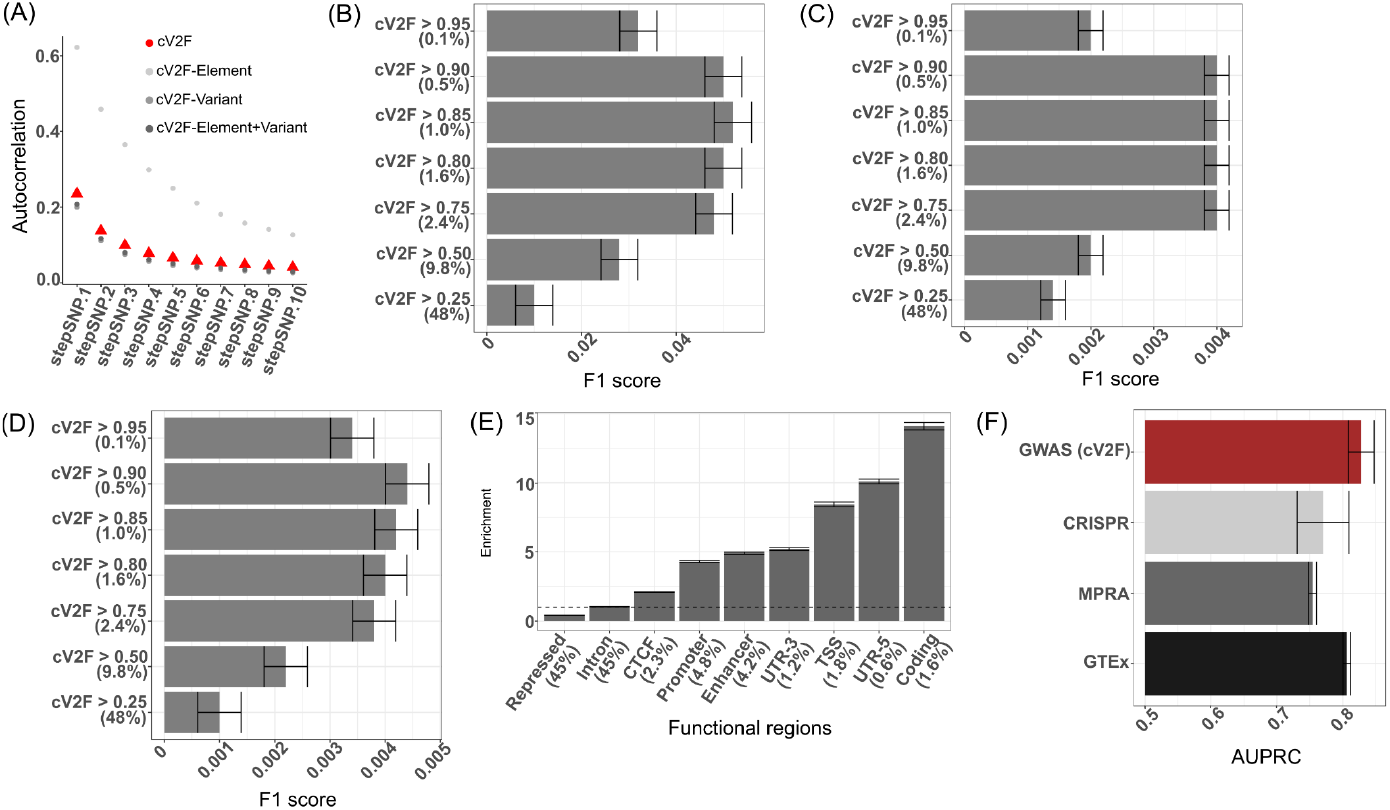
(A) The autocorrelation in the primary binarized cV2F scores and secondary binarized cV2F scores created by taking different broad subsets of features, for variants that are between 1st and 10th nearest in physical distance to each variant in the genome. (Panels B-D) F1 score (geometric mean of precision and recall) of variants annotated by cV2F at different probability thresholds when evaluated against (B) MPRA positive versus tested variants, (C) WG-STARR-seq positive versus tested variants, and (D) GWAS confidently fine-mapped variants (PIP> 0.90) from 94 UKBB traits. (E) The excess overlap of the binarized cV2F scores (cV2F thresholded at 0.75) with respect to different functional regions of the genome such enhancers, promoters, coding regions etc. Error bars denote 95% confidence intervals. (F) Area under the Precision Recall Curve (AUPRC) on held-out chromosome data for gradient boosting model on all 339 cV2F features but using training data from CRISPR, MPRA and GTEx finemapped eQTLs. Error bars represent 95% confidence intervals. Numerical results are reported in **Supplementary Table S2**.

**Supplementary Figure S3:**
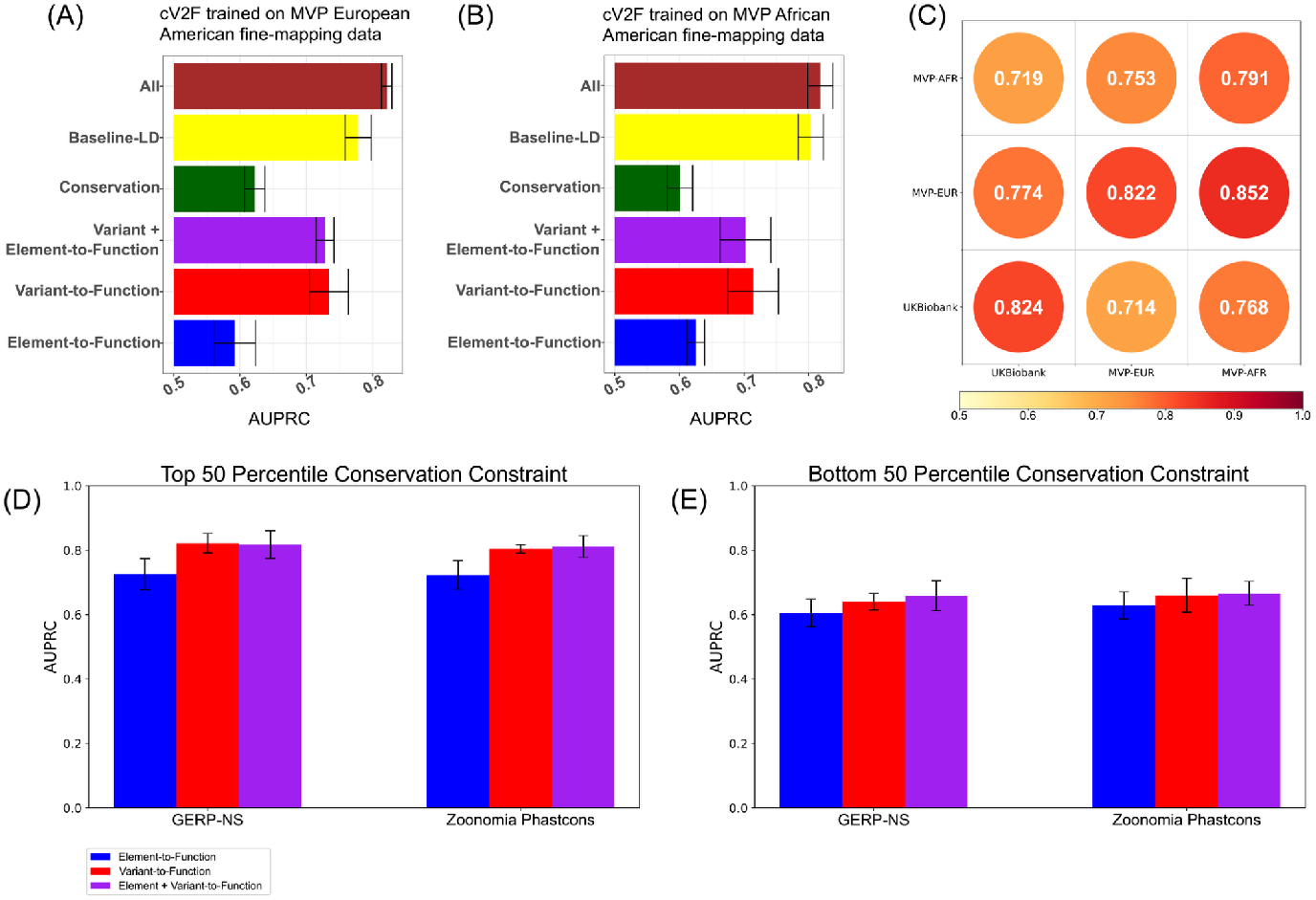
Area under the Precision Recall curve (AUPRC) on held-out chromosomes for the several broad categories of functional features when using 931 Million Veteran Program GWAS traits for training instead of the 94 UKBiobank traits used in the primary analysis. We considered the positive set to be PIP> 0.90 variants in at least one trait based on ancestry-specific statistical fine-mapping of GWAS signals in (A) European-Americans and (B) African-Americans. The negative set of variants were selected by variants with low PIP (PIP < 0.01) across all traits that are LD and MAF matched to the positive set using European and African LD reference panels^117^ (in the absence of in-sample LD reference panels from MVP). We considered 4 broad categories of functional features - all 339 features, 84 baseline-LD features, 17 new conservation features from Zoonomia and FUNCODE, 238 element and variant-level function, 186 variant-level functional features, and 52 element-level functional features. Error bars denote 95% confidence intervals. (C) Test AUPRC observed when the 339 cV2F features are separately trained on fine-mapping data from UKBiobank (Europeans), MVP (European Americans), and MVP (African Americans), and tested on held out chromosome data from all three cohorts. (D, E) AUPRC of element-level, variant-level and element plus variant-level features on UKBB training data subsetted into two bins of GERP and Zoonomia PhastCons scores - the results for the training data in the top (respectively bottom) 50 percentile of either score is reported in panel D (respectively panel E). Numerical results are reported in **Supplementary Table S2**.

**Supplementary Figure S4:**
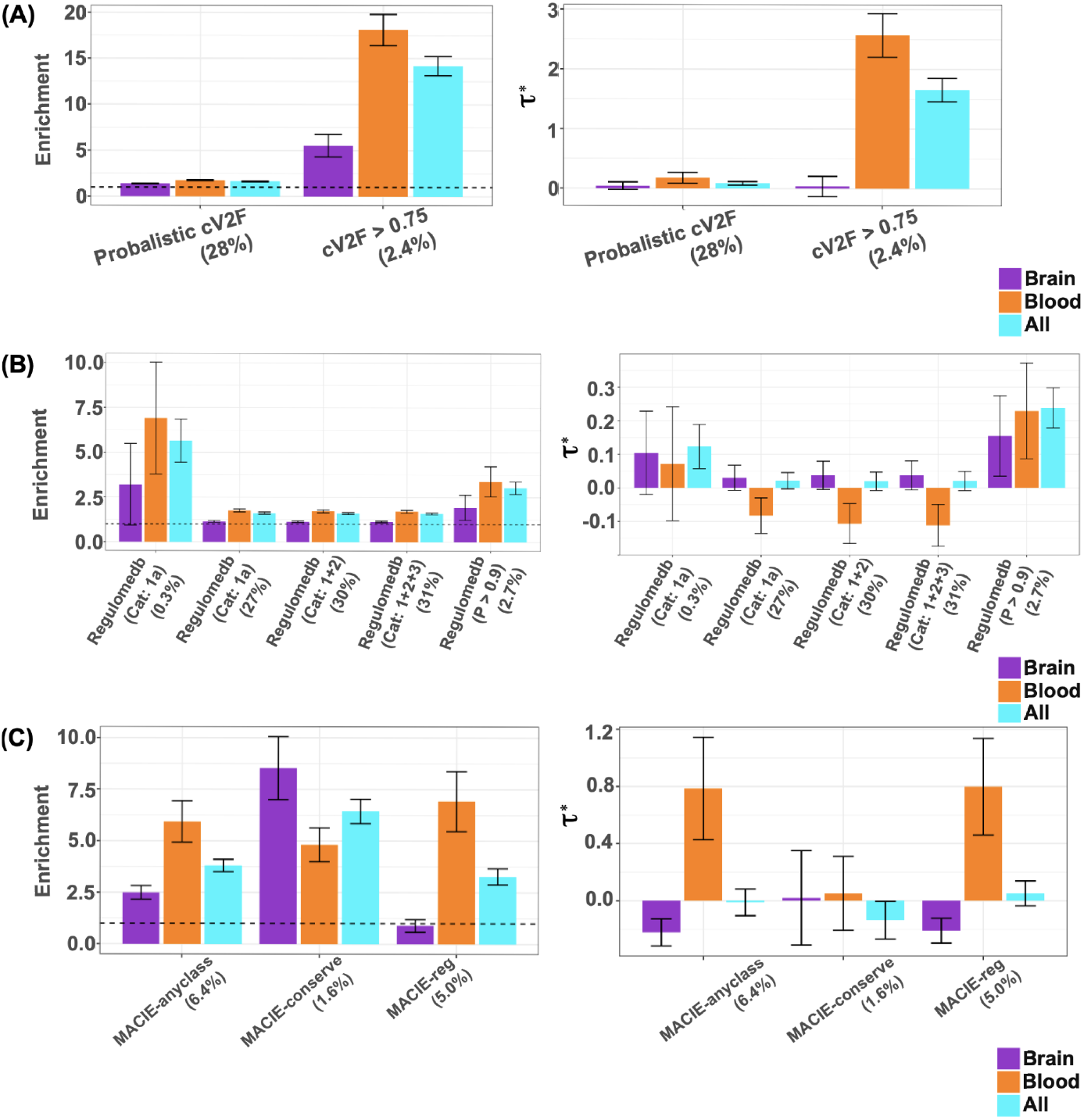
S-LDSC Heritability enrichment (left) and standardized effect sizes (***τ***\*) (right) of (A) probabilistic and binarized cV2F score, (B) 5 different categories of Regulomedb (v2.2) scores, and (C) 3 broad categories of MACIE scores. Results are conditional on 97 baseline-LD v2.2 annotations. Dashed horizontal line denotes no enrichment. Results are meta-analyzed across all 66 relatively independent diseases and traits, as well as 15 relatively independent blood-related traits and 10 relatively independent brain-related traits. Error bars denote 95% confidence intervals. Numerical results are reported in **Supplementary Table S5**.

**Supplementary Figure S5:**
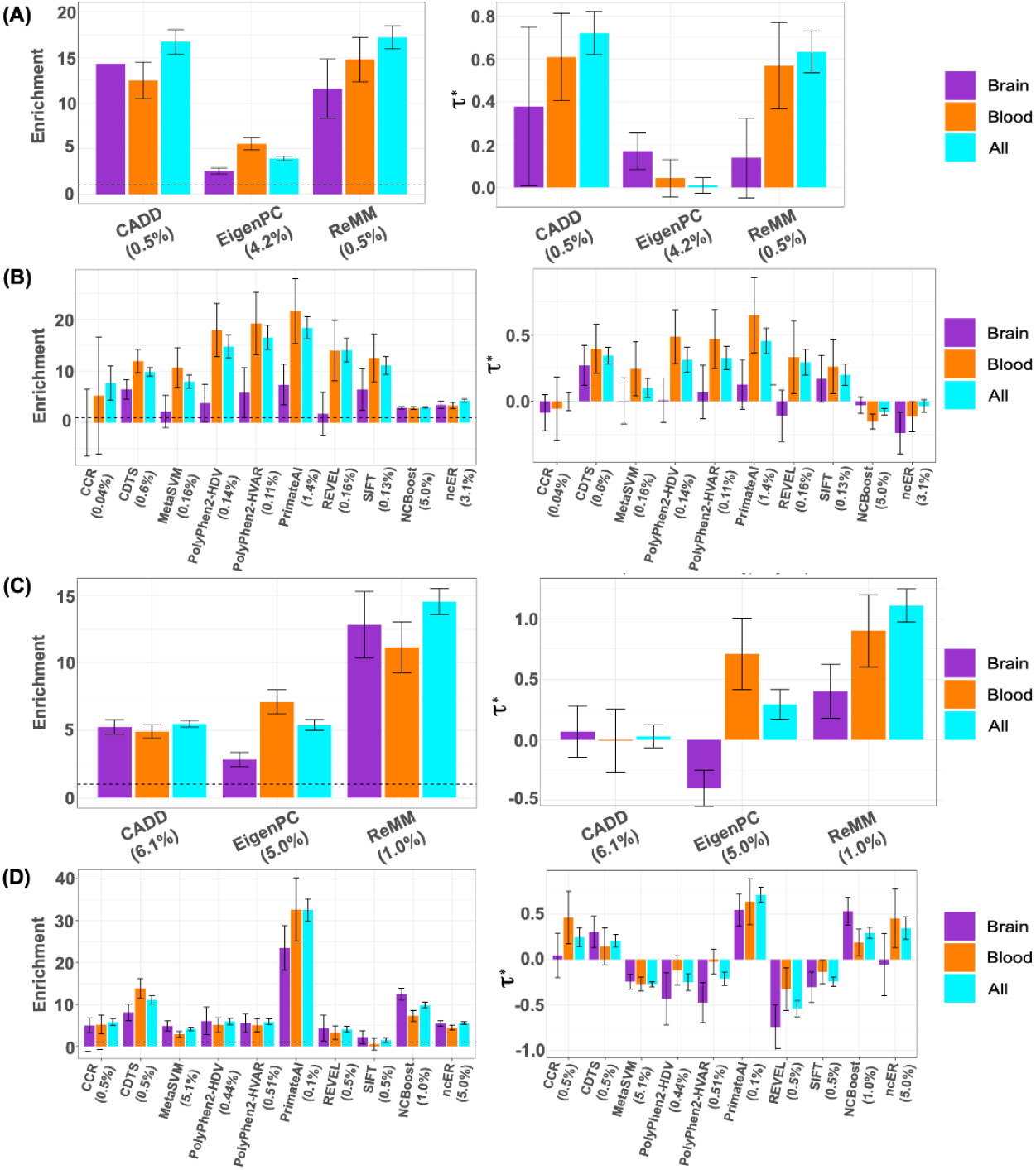
S-LDSC Heritability enrichment (left) and standardized effect sizes (***τ***\*) (right) of (A) 3 primary pathogenicity scores from Figure 3, (B) 10 secondary pathogenicity scores, (C) AnnotBoost boosted genome-wide versions^24^ of 3 primary pathogenicity scores, and (D) AnnotBoost boosted versions of 10 secondary pathogenicity scores. Results are conditional on 97 baseline-LD annotations. Dashed horizontal line denotes no enrichment. Results are meta-analyzed across all 66 relatively independent diseases and traits, as well as 15 relatively independent blood-related traits and 10 relatively independent brain-related traits. Error bars denote 95% confidence intervals. Numerical results are reported in **Supplementary Table S5**.

**Supplementary Figure S6:**
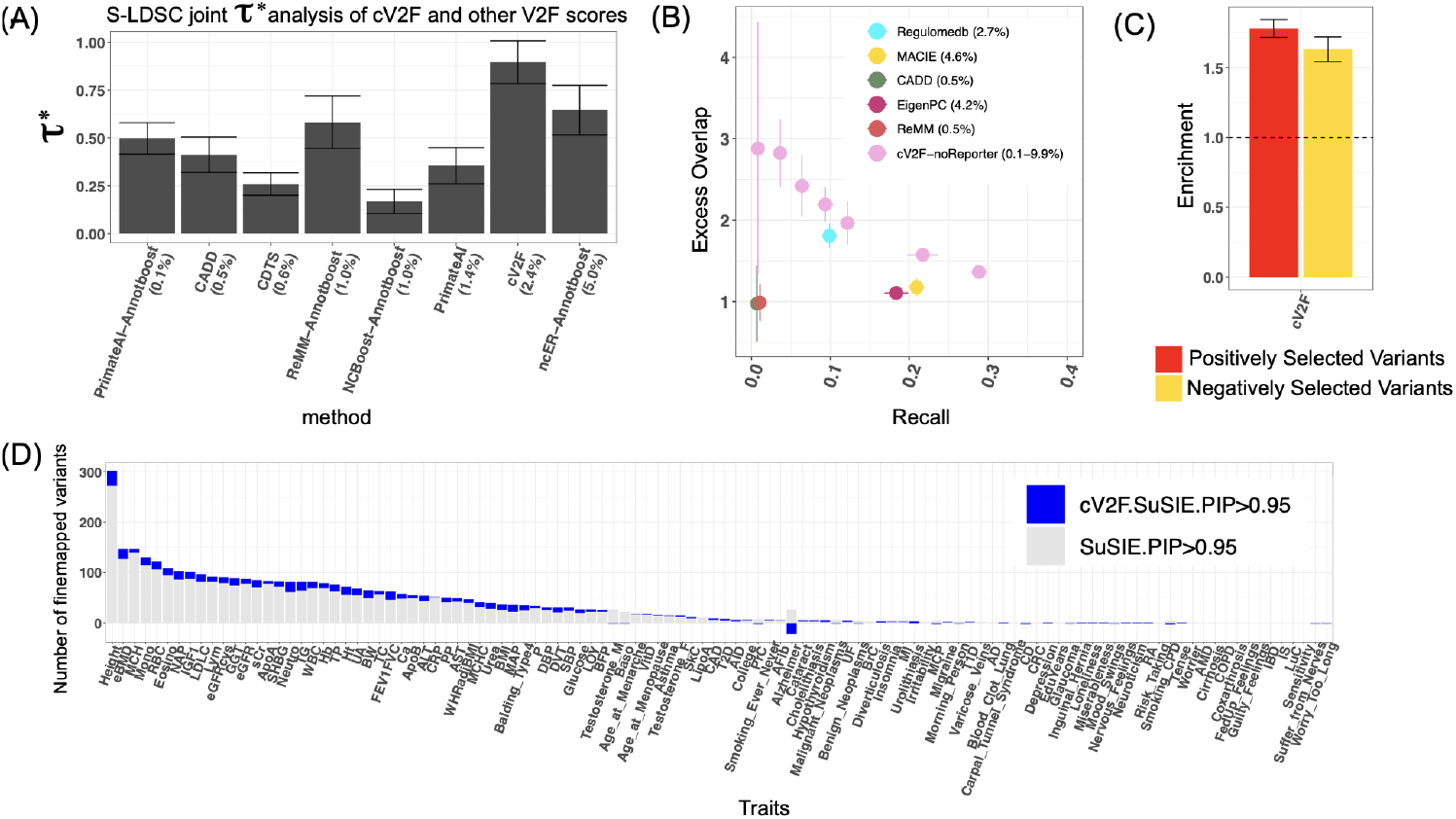
(A) S-LDSC standardized effect sizes (τ*) (bottom panel) of 8 jointly significant annotations in a joint model comprising of binarized cV2F, MACIE, Regulomedb, primary and secondary pathogenicity scores and their boosted versions, and the baseline-LD v2.2 model annotations. Annotations are displayed in order of their annotation sizes from left to right. Results are meta-analyzed across all 66 relatively independent diseases and traits. (B) Excess overlap and recall of variants annotated by different variant-function and pathogenicity scores with respect to variants that are Whole Genome STARR-seq positives among the ones tested for functional characterization, in comparison to the cV2F-noReporter score (an analogous score to cV2F but not including any MPRA related features), binarized at different thresholds. (C) Excess overlap of variants annotated by optimally binarized cV2F in 21,129 candidate positively selected and 24,152 candidate negatively selected variants based on genetic adaptation to selective pressure exerted by pathogens^78^. (D) Number of variants confidently fine-mapped (posterior probability of causality > 0.95) from standard fine-mapping and cV2F-informed functional fine-mapping of 94 UKBB diseases and traits. Error bars denote 95% confidence intervals. Numerical results are reported in **Supplementary Tables S4, S5, S6**.

**Supplementary Figure S7:**
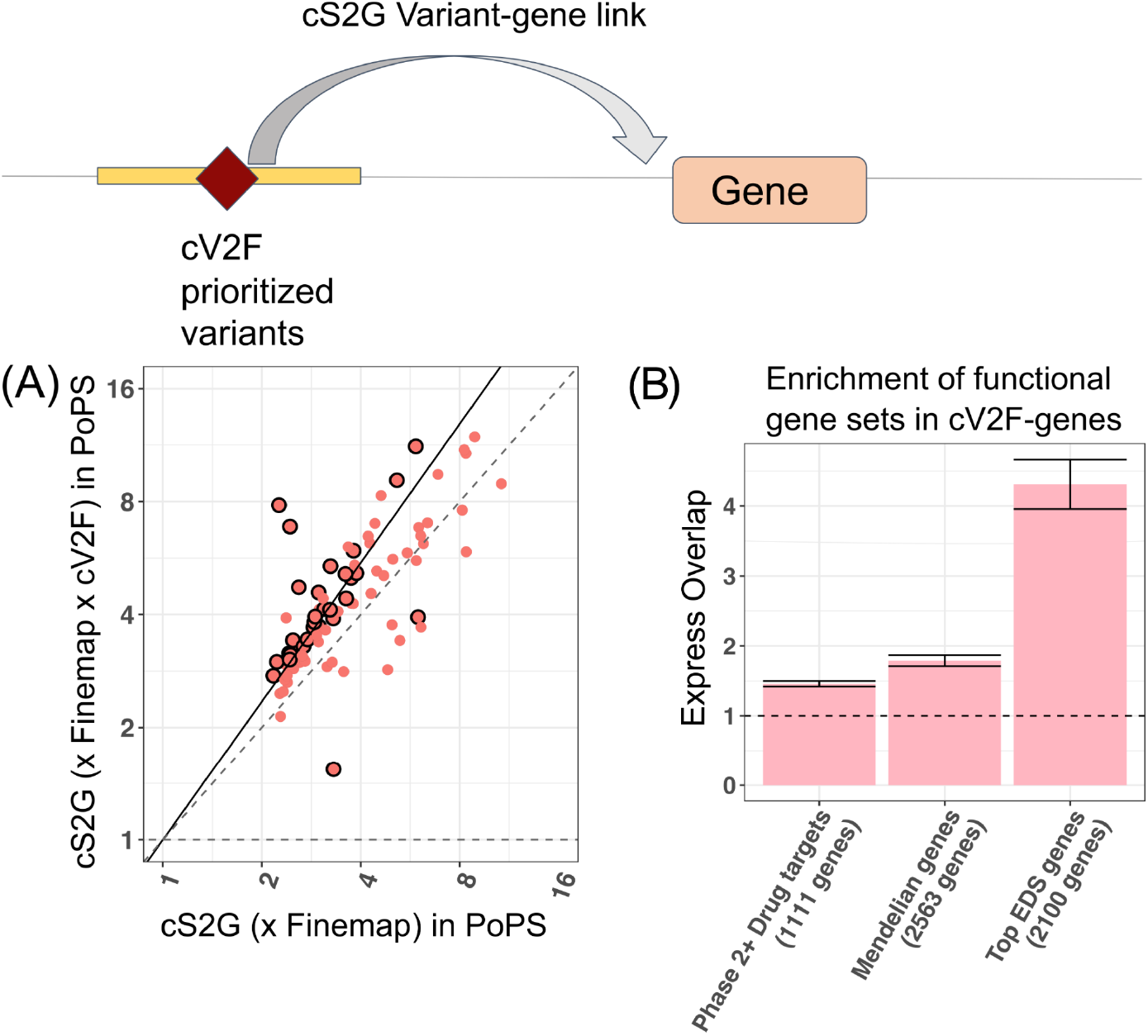
(A) Average PoPS disease prioritization scores^23^ of genes linked to weakly fine-mapped variants (PIP > 0.10) by the combined S2G (cS2G)^79^ with and without restricting to variants annotated by the optimally binarized cV2F (thresholded at 0.75). Each point corresponds to one of 94 UKBiobank diseases and traits. Circled dots represent traits for which we see significant (FDR < 10%) difference in average PoPs scores for genes linked to weakly fine-mapped variants and genes linked to weakly fine-mapped variants that are also implicated by cV2F. The solid line denotes y=x, and the dashed line denotes the regression slope. We report the slope and the p-value of the regression coefficient from the regression model. (B) Excess overlap of top 10% genes with the most number of binarized cV2F annotated variants linked using the cS2G approach, with respect to genes that are approved drug targets^80^, genes linked to Mendelian disorders^81^ and genes in top 10% of the Enhancer Domain Score (EDS)^82^. Error bars denote 95% confidence intervals. Dashed horizontal line denotes no excess overlap. Numerical results are reported in **Supplementary Table S4**.

**Supplementary Figure S8:**
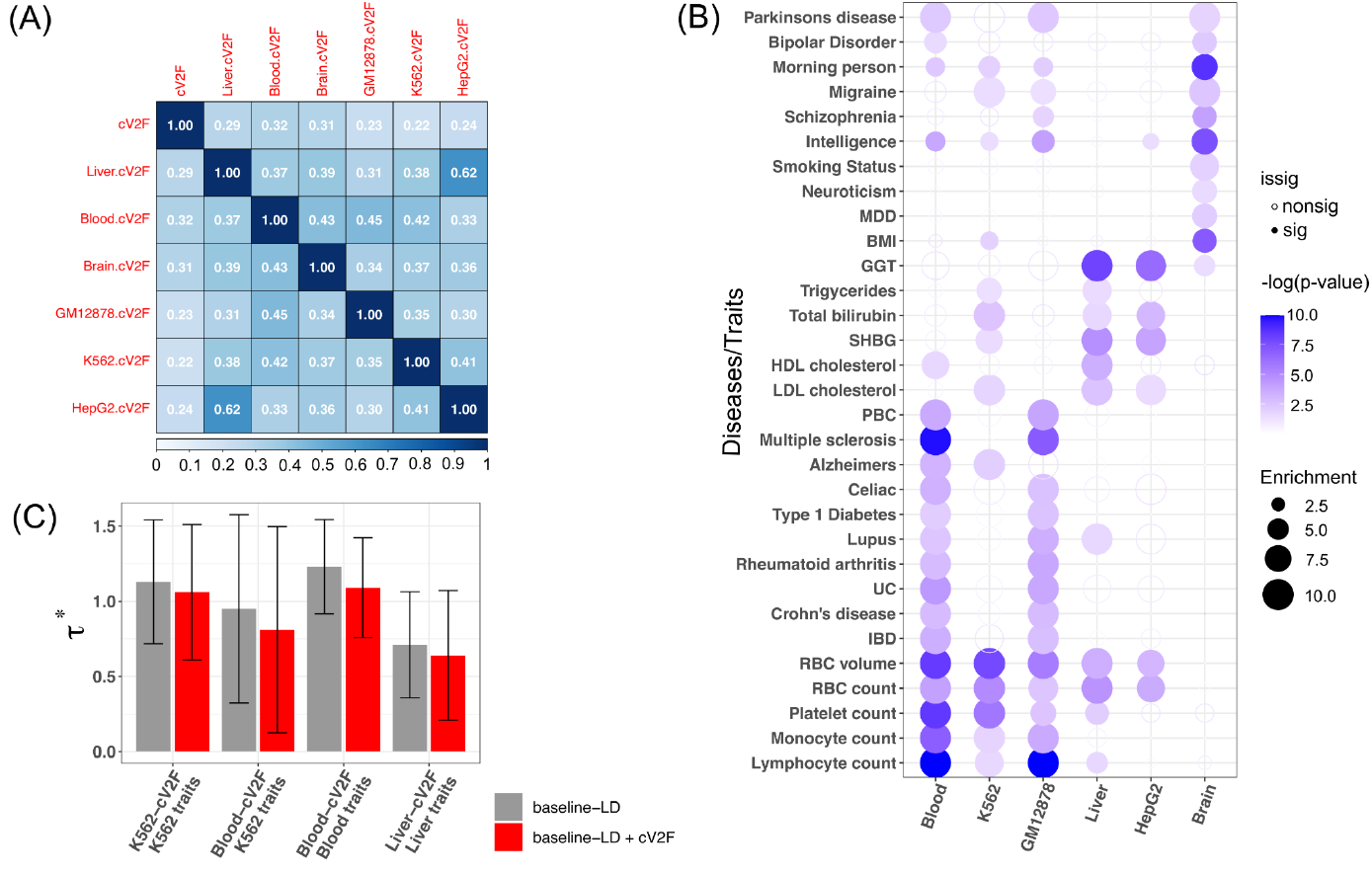
(A) Correlation of binarized primary and cell-line and tissue-specific cV2F scores. (B) For specific cell-line or tissue-specific cV2F, meta-analyzed S-LDSC standardized effect sizes corresponding to a set of related diseases and traits, either conditional on the 97 baseline-LD model annotations (colored gray), or 97 baseline-LD + 1 standard binarized cV2F annotation (colored red). Error bars denote 95% confidence intervals. (C) S-LDSC heritability enrichment of cell-line and tissue-specific cV2F scores for a set of related diseases and traits. Results are conditional on 97 baseline-LD v2.2 + 1 binarized primary cV2F annotations. Magnitude (Enrichment, dot size) and significance (−log10(*P*), dot color) are reported for disease signal for 31 blood, liver and brain-related traits. Numerical results are reported in **Supplementary Table S7**.

**Supplementary Figure S9:**
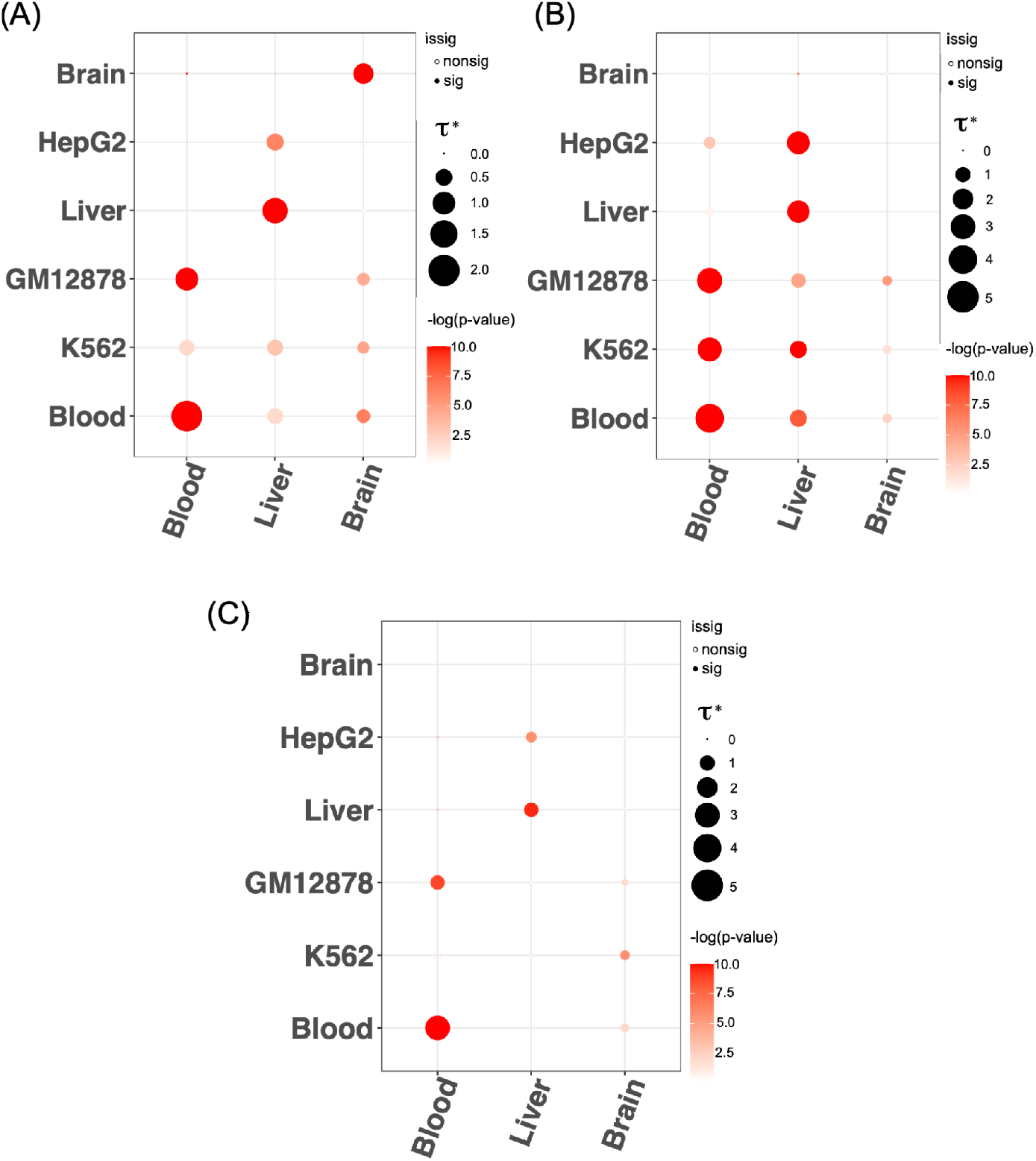
(A) S-LDSC meta-analyzed standardized effect sizes (τ*) of 6 cell-line and tissue-specific cV2F scores (along Y-axis) across blood-related, liver-related and brain-related traits. (B) Same analysis as (A) but performed for secondary cell-line and tissue-specific cV2F scores obtained using a gradient boosting model trained using all 339 features on GWAS fine-mapping data specific to blood, liver and brain-related traits. (C) Same analysis as (A) but performed for secondary cell-line and tissue-specific cV2F scores obtained using a gradient boosting model trained on tissue-specific GWAS fine-mapping training data and tissue-specific features. Magnitude (τ*, dot size) and significance (−log10(*P*), dot color) of the standardized effect sizes of cell-line and tissue-specific cV2F scores. Numerical results are reported in **Supplementary Table S7**.

