## Supplementary Figures for "A consensus variant-to-function score to functionally prioritize variants for disease"

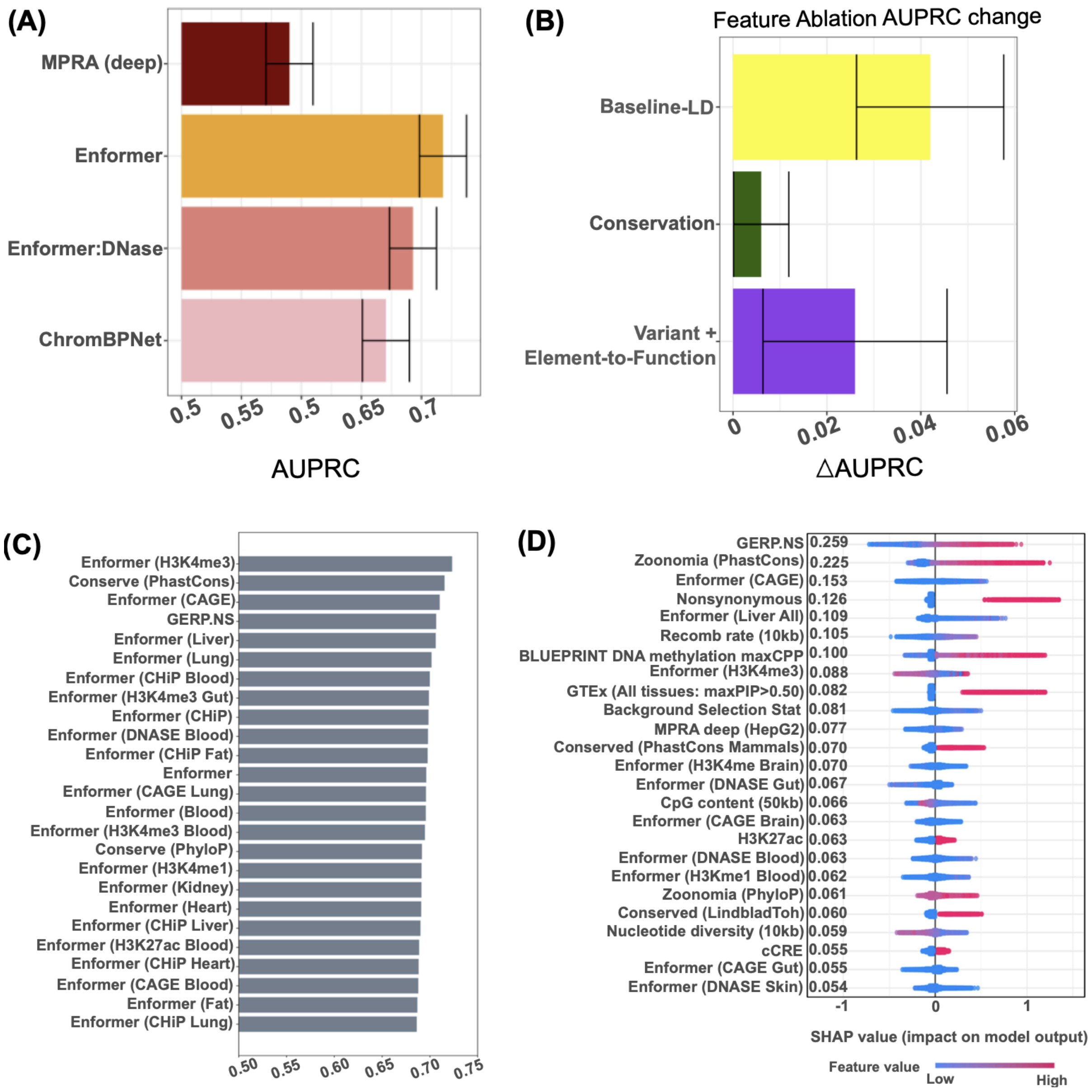


**Supplementary Figure S1:** (A) The AUPRC of the GWAS fine-mapping gradient boosting training model corresponding to deep learning predicted allelic effect functional annotations from MPRA-based deep learning model, Enformer, Enformer restricted to DNase features and ChromBPNet features. (B) Change in Area Under the Precision Recall Curve (AUPRC) of the cV2F GWAS fine-mapping based gradient boosting training model upon ablation of three broad categories of functional annotations - baseline-LD, conservation annotations (Zoonomia and FUNCODE) and ENCODE element and variant-level functional annotations. (C) The AUPRC of the gradient boosting training model for individual functional features; results are reported for the top 25 features. See Supplementary Table 2 for results from the full set of 339 features. (D) The Shapley values of the top 25 predictive features based on the cV2F gradient boosting training model. Error bars denote 95% confidence intervals. Numerical results are reported in **Supplementary Table S2**.

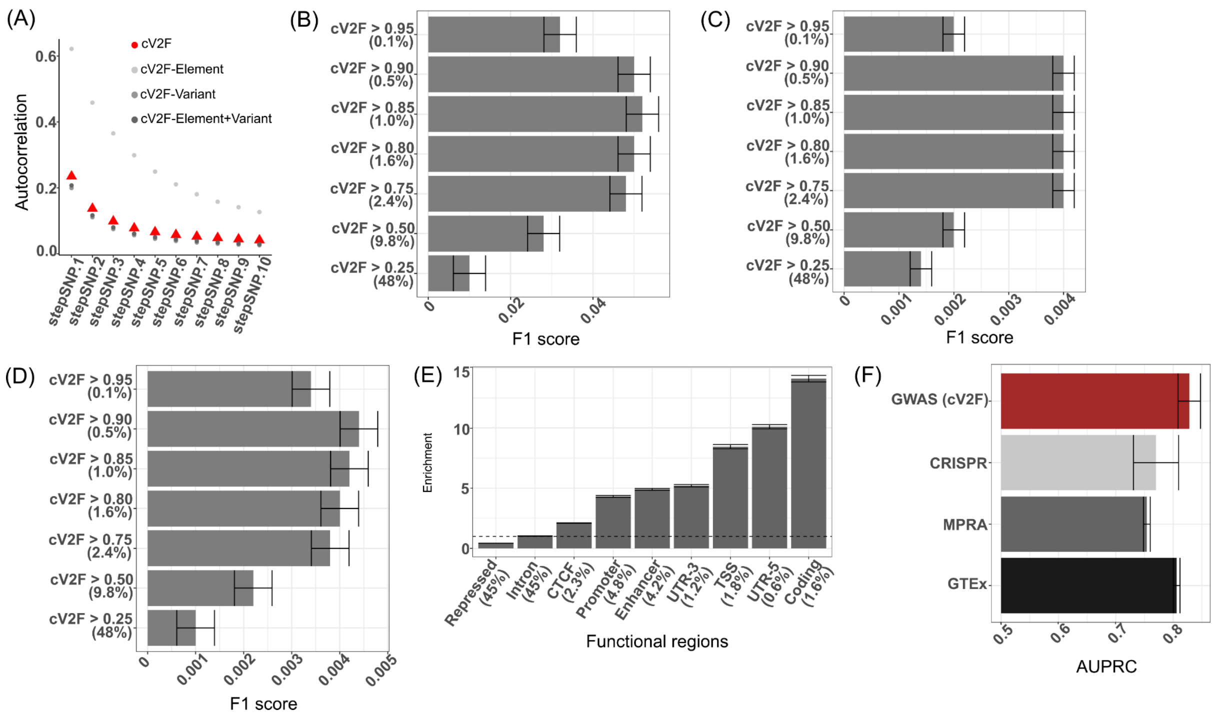


**Supplementary Figure S2:** (A) The autocorrelation in the primary binarized cV2F scores and secondary binarized cV2F scores created by taking different broad subsets of features, for variants that are between 1st and 10th nearest in physical distance to each variant in the genome. **(**Panels B-D**)** F1 score (geometric mean of precision and recall) of variants annotated by cV2F at different probability thresholds when evaluated against (B) MPRA positive versus tested variants, (C) WG-STARR-seq positive versus tested variants, and (D) GWAS confidently fine-mapped variants (PIP> 0.90) from 94 UKBB traits. (E) The excess overlap of the binarized cV2F scores (cV2F thresholded at 0.75) with respect to different functional regions of the genome such enhancers, promoters, coding regions etc. Error bars denote 95% confidence intervals. (F) Area under the Precision Recall Curve (AUPRC) on held-out chromosome data for gradient boosting model on all 339 cV2F features but using training data from CRISPR, MPRA and GTEx finemapped eQTLs. Error bars represent 95% confidence intervals. Numerical results are reported in **Supplementary Table S2**.

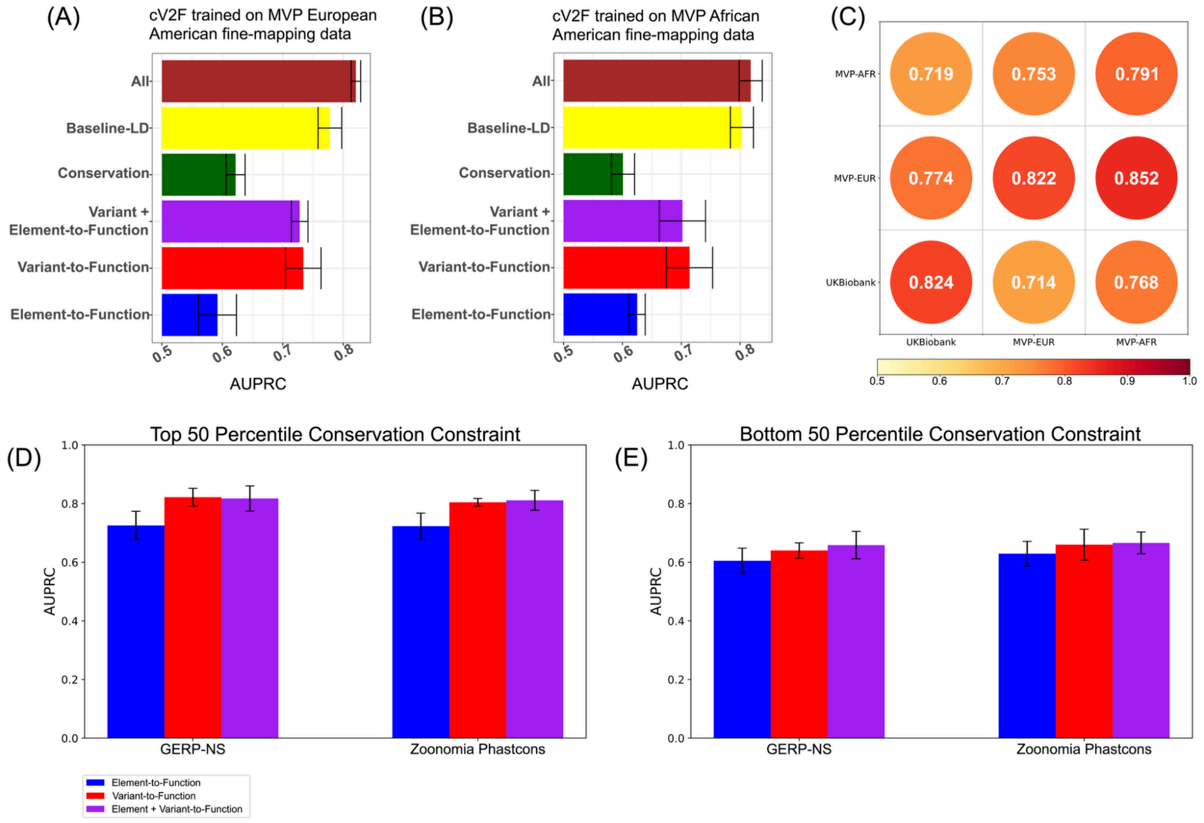


**Supplementary Figure S3:** Area under the Precision Recall curve (AUPRC) on held-out chromosomes for the several broad categories of functional features when using 931 Million Veteran Program GWAS traits for training instead of the 94 UKBiobank traits used in the primary analysis. We considered the positive set to be PIP> 0.90 variants in at least one trait based on ancestry-specific statistical fine-mapping of GWAS signals in (A) European-Americans and (B) African-Americans. The negative set of variants were selected by variants with low PIP (PIP < 0.01) across all traits that are LD and MAF matched to the positive set using European and African LD reference panels[117] (in the absence of in-sample LD reference panels from MVP). We considered 4 broad categories of functional features - all 339 features, 84 baseline-LD features, 17 new conservation features from Zoonomia and FUNCODE, 238 element and variant-level function, 186 variant-level functional features, and 52 element-level functional features. Error bars denote 95% confidence intervals. (C ) Test AUPRC observed when the 339 cV2F features are separately trained on fine-mapping data from UKBiobank (Europeans), MVP (European Americans), and MVP (African Americans), and tested on held out chromosome data from all three cohorts. (D, E) AUPRC of element-level, variant-level and element plus variant-level features on UKBB training data subsetted into two bins of GERP and Zoonomia PhastCons scores - the results for the training data in the top (respectively bottom) 50 percentile of either score is reported in panel D (respectively panel E). Numerical results are reported in **Supplementary Table S2**.

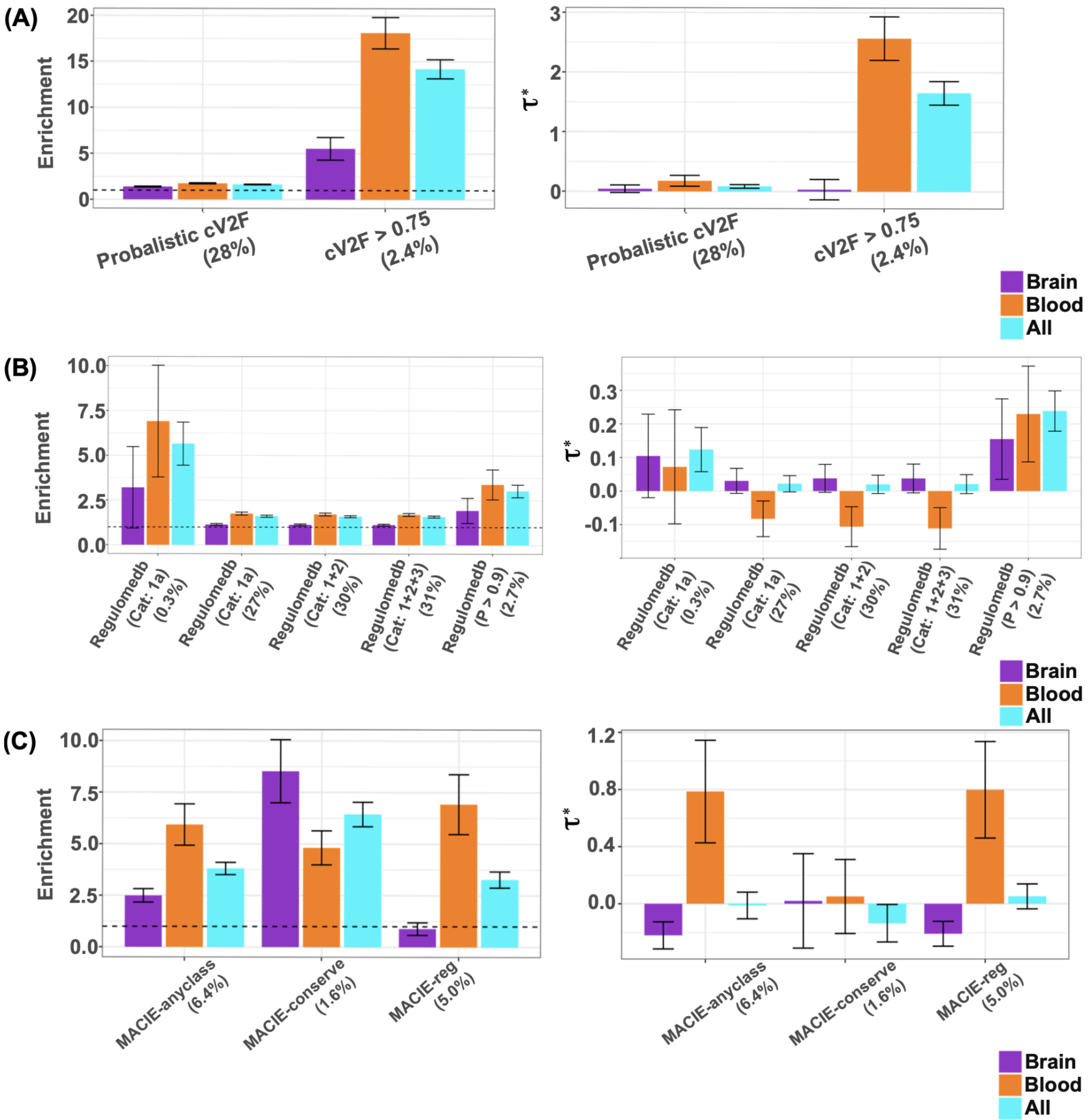


**Supplementary Figure S4:** S-LDSC Heritability enrichment (left) and standardized effect sizes (**τ***) (right) of (A) probabilistic and binarized cV2F score, (B) 5 different categories of Regulomedb (v2.2) scores, and (C ) 3 broad categories of MACIE scores. Results are conditional on 97 baseline-LD v2.2 annotations. Dashed horizontal line denotes no enrichment. Results are meta-analyzed across all 66 relatively independent diseases and traits, as well as 15 relatively independent blood-related traits and 10 relatively independent brain-related traits. Error bars denote 95% confidence intervals. Numerical results are reported in **Supplementary Table S5**.

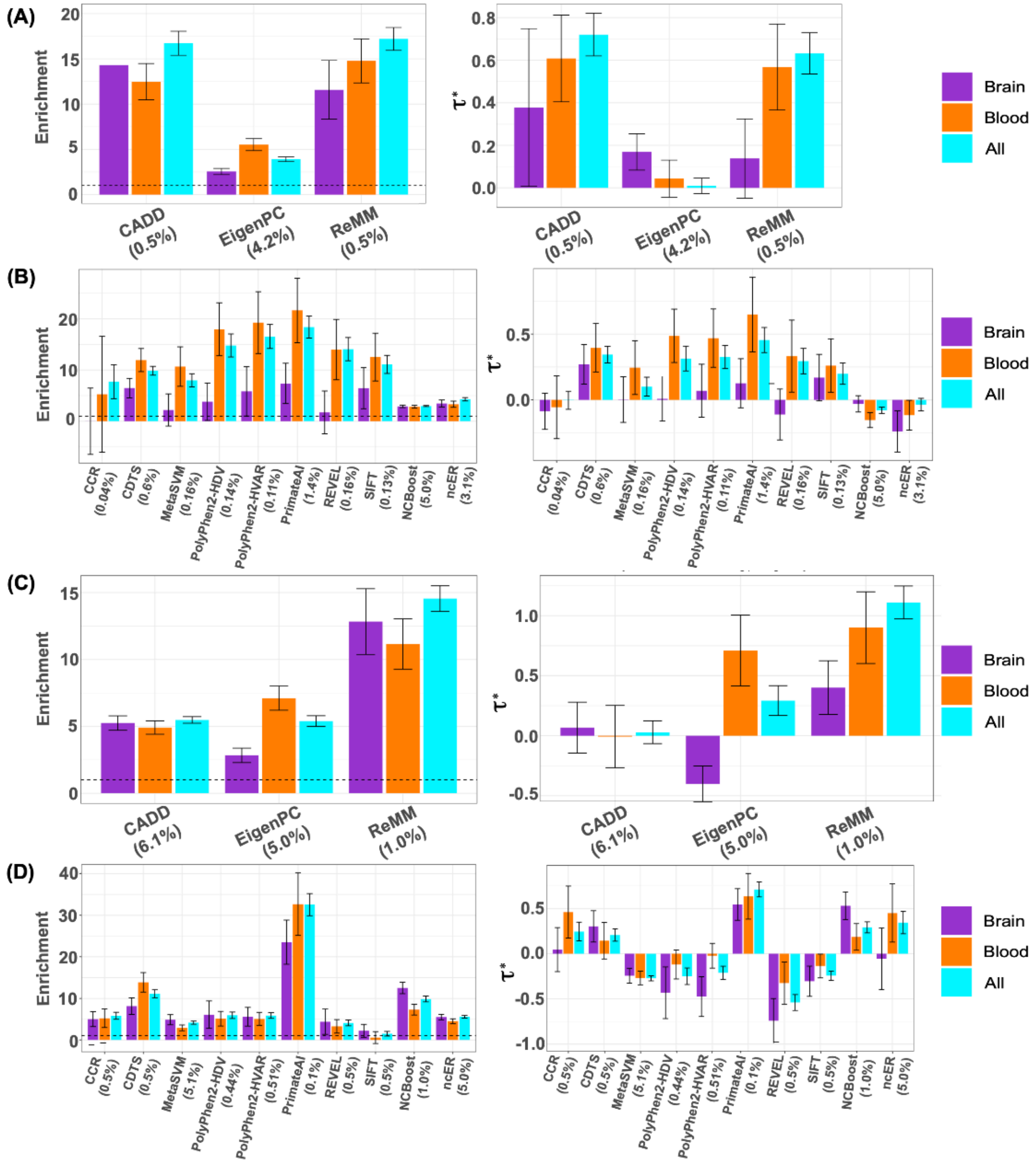


**Supplementary Figure S5:** S-LDSC Heritability enrichment (left) and standardized effect sizes (**τ***) (right) of (A) 3 primary pathogenicity scores from Figure 3 , (B) 10 secondary pathogenicity scores,  (C ) AnnotBoost boosted genome-wide versions[24] of 3 primary pathogenicity scores, and (D) AnnotBoost boosted versions of 10 secondary pathogenicity scores. Results are conditional on 97 baseline-LD  annotations. Dashed horizontal line denotes no enrichment. Results are meta-analyzed across all 66 relatively independent diseases and traits, as well as 15 relatively independent blood-related traits and 10 relatively independent brain-related traits. Error bars denote 95% confidence intervals. Numerical results are reported in **Supplementary Table S5**.

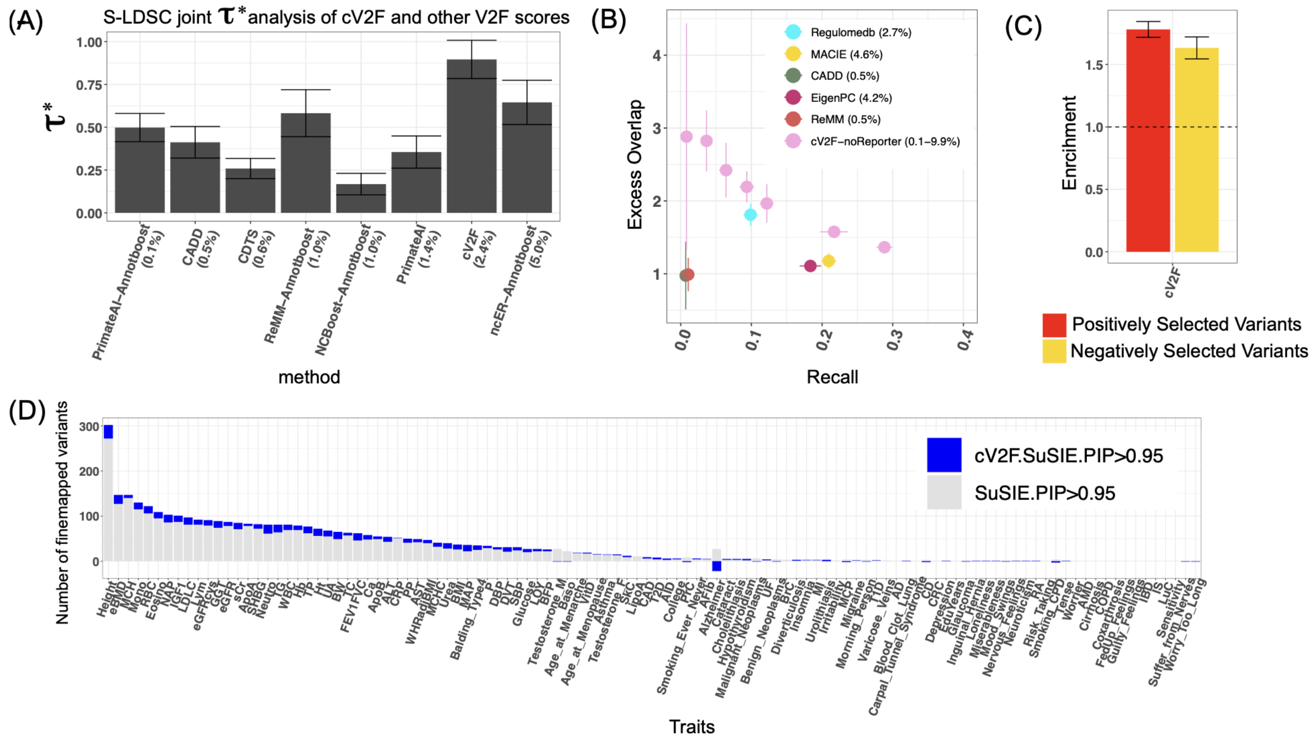


**Supplementary Figure S6:** (A) S-LDSC standardized effect sizes (τ*) (bottom panel) of 8 jointly significant annotations in a joint model comprising of binarized cV2F, MACIE, Regulomedb, primary and secondary pathogenicity scores and their boosted versions, and the baseline-LD v2.2 model annotations. Annotations are displayed in order of their annotation sizes from left to right. Results are meta-analyzed across all 66 relatively independent diseases and traits. (B) Excess overlap and recall of variants annotated by different variant-function and pathogenicity scores with respect to variants that are Whole Genome STARR-seq positives among the ones tested for functional characterization, in comparison to the cV2F-noReporter score (an analogous score to cV2F but not including any MPRA related features), binarized at different thresholds. (C) Excess overlap of variants annotated by optimally binarized cV2F in 21,129 candidate positively selected and 24,152 candidate negatively selected variants based on genetic adaptation to selective pressure exerted by pathogens[78]. (D) Number of variants confidently fine-mapped (posterior probability of causality > 0.95) from standard fine-mapping and cV2F-informed functional fine-mapping of 94 UKBB diseases and traits. Error bars denote 95% confidence intervals. Numerical results are reported in **Supplementary Tables S4, S5, S6**.

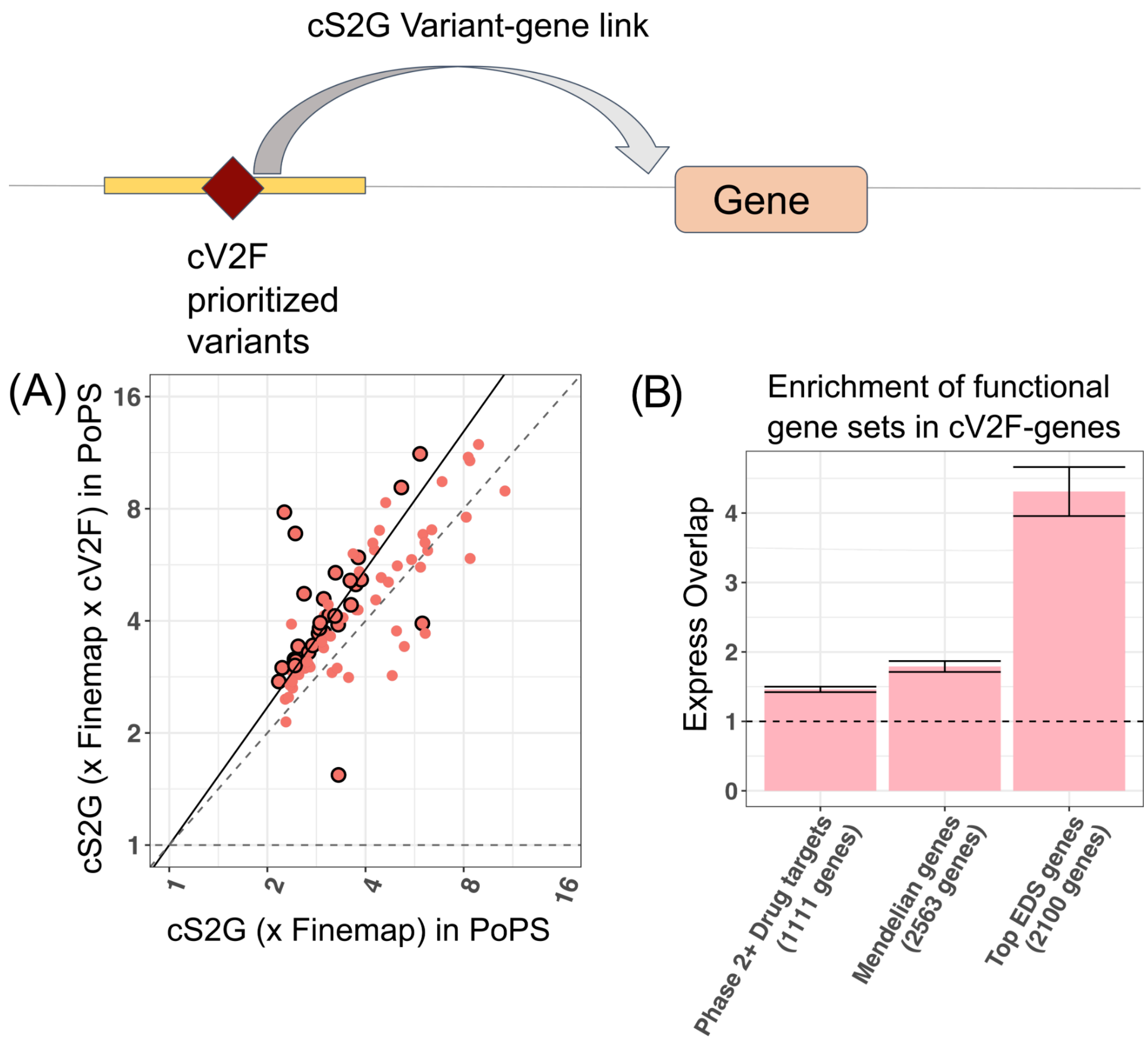


**Supplementary Figure S7:** (A) Average PoPS disease prioritization scores[23] of genes linked to weakly fine-mapped variants (PIP > 0.10) by the combined S2G (cS2G)[79] with and without restricting to variants annotated by the optimally binarized cV2F (thresholded at 0.75). Each point corresponds to one of 94 UKBiobank diseases and traits. Circled dots represent traits for which we see significant (FDR < 10%) difference in average PoPs scores for genes linked to weakly fine-mapped variants and genes linked to weakly fine-mapped variants that are also implicated by cV2F. The solid line denotes y=x, and the dashed line denotes the regression slope. We report the slope and the p-value of the regression coefficient from the regression model. (B) Excess overlap of top 10% genes with the most number of binarized cV2F annotated variants linked using the cS2G approach, with respect to genes that are approved drug targets[80], genes linked to Mendelian disorders[81] and genes in top 10% of the Enhancer Domain Score (EDS)[82]. Error bars denote 95% confidence intervals. Dashed horizontal line denotes no excess overlap. Numerical results are reported in **Supplementary Table S4**.

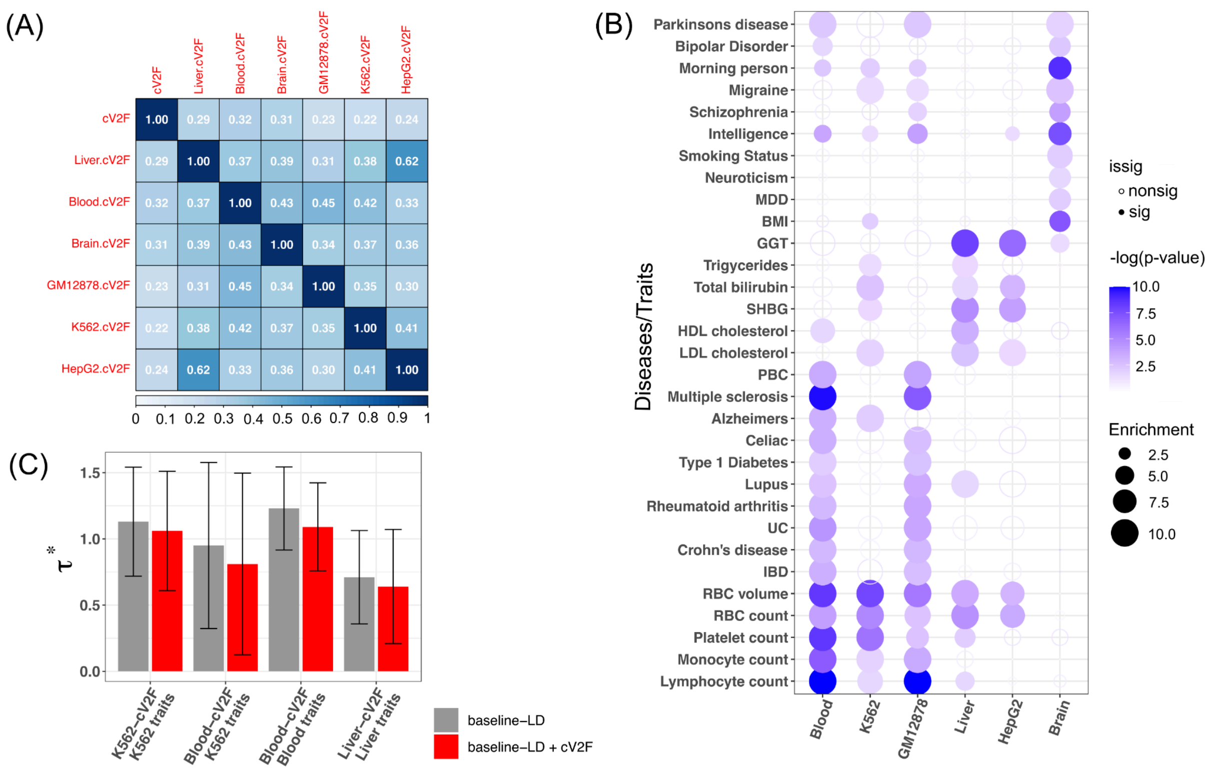


**Supplementary Figure S8:** (A) Correlation of binarized primary and cell-line and tissue-specific cV2F scores. (B) For specific cell-line or tissue-specific cV2F, meta-analyzed S-LDSC standardized effect sizes corresponding to a set of related diseases and traits, either conditional on the 97 baseline-LD model annotations (colored gray), or 97 baseline-LD + 1 standard binarized cV2F annotation (colored red). Error bars denote 95% confidence intervals. (C) S-LDSC heritability enrichment of cell-line and tissue-specific cV2F scores for a set of related diseases and traits. Results are conditional on 97 baseline-LD v2.2 + 1 binarized primary cV2F annotations. Magnitude (Enrichment, dot size) and significance (−log10($P$), dot color) are reported for disease signal for 31 blood, liver and brain-related traits. Numerical results are reported in **Supplementary Table S7**.

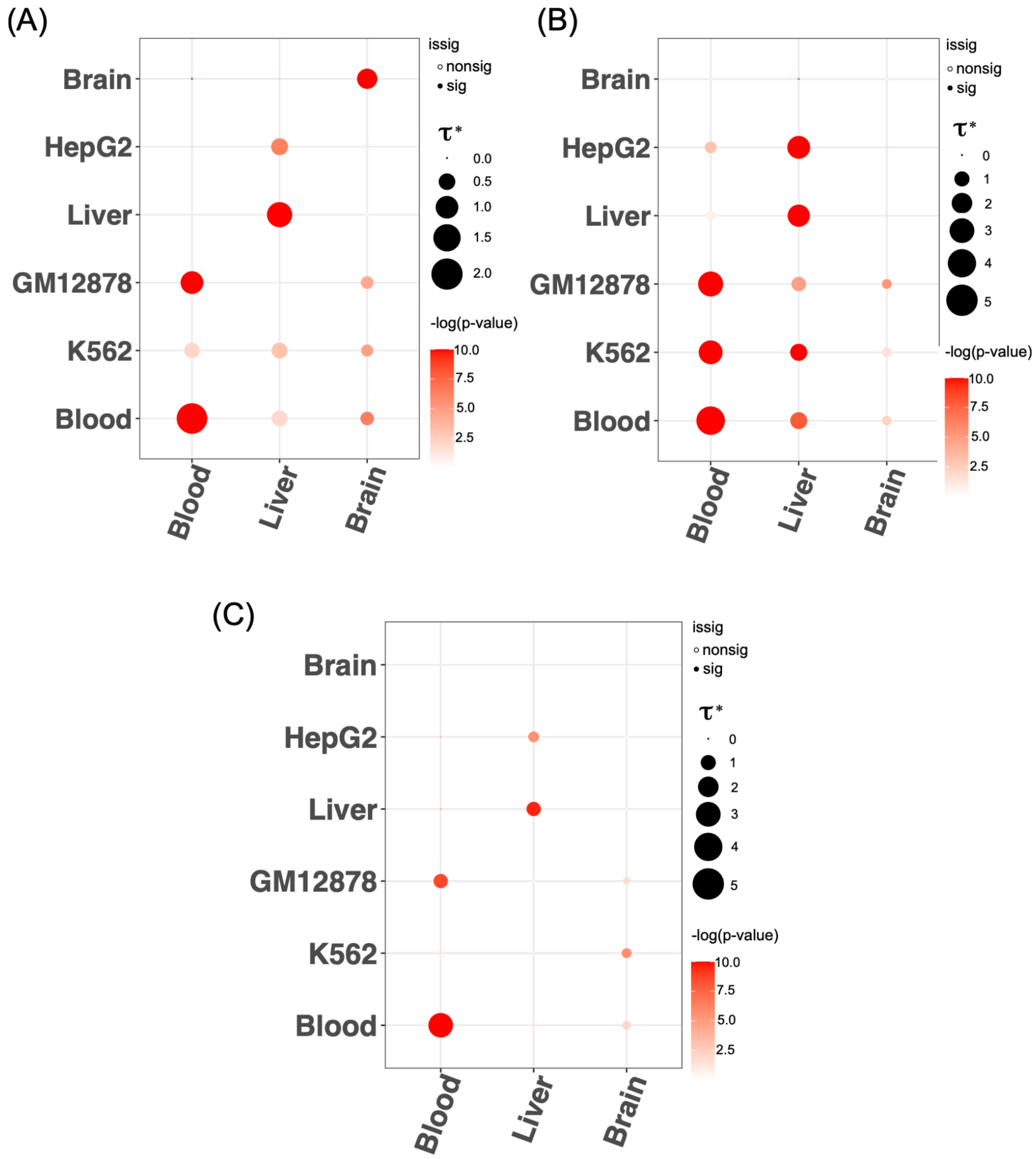


**Supplementary Figure S9:** (A) S-LDSC meta-analyzed standardized effect sizes (τ*) of 6 cell-line and tissue-specific cV2F scores (along Y-axis) across blood-related, liver-related and brain-related traits. (B) Same analysis as (A) but performed for secondary cell-line and tissue-specific cV2F scores obtained using a gradient boosting model trained using all 339 features on GWAS fine-mapping data specific to blood, liver and brain-related traits. (C) Same analysis as (A) but performed for secondary cell-line and tissue-specific cV2F scores obtained using a gradient boosting model trained on tissue-specific GWAS fine-mapping training data and tissue-specific features. Magnitude (τ*, dot size) and significance (−log10(*P*), dot color) of the standardized effect sizes of cell-line and tissue-specific cV2F scores. Numerical results are reported in **Supplementary Table S7**.
